# Architecting *cis*-regulation to quantitatively tune gene expression in cereals

**DOI:** 10.64898/2026.01.08.698005

**Authors:** Evan D. Groover, David Ding, Flora Z. Wang, Gonzalo Benegas, Joseph Rivera, Shahar Schwartz, Stephen Chen, Michael Moubarak, Viktoriya Georgieva, Peggy G. Lemaux, Brian J. Staskawicz, Krishna K. Niyogi, Yun S. Song, David F. Savage

**Affiliations:** Innovative Genomics Institute, University of California, Berkeley, CA 94720, USA; Department of Plant and Microbial Biology, University of California, Berkeley, CA 94720, USA; Department of Molecular and Cellular Biology, University of California, Berkeley, CA 94720, USA; Howard Hughes Medical Institute, University of California, Berkeley, CA 94720, USA; Department of Electrical Engineering and Computer Sciences, University of California, Berkeley, CA, US; Center for Computational Biology, University of California, Berkeley, CA 94720, USA; Molecular Biophysics and Integrated Bioimaging Division, Lawrence Berkeley National Laboratory, Berkeley, CA, 94720 USA; Department of Statistics, University of California, Berkeley, CA, USA

## Abstract

Precise modulation of gene expression via *cis*-regulatory editing holds promise for non-transgenic crop improvement, but the sequence-to-function relationships that govern plant promoter activity remain poorly understood. Here, we develop a massively parallel reporter assay (MPRA) in *Sorghum bicolor* to systematically measure the effects of >30,000 CRISPR-like mutations-deletions, substitutions, and motif insertions-across entire native promoters and 5′ untranslated regions (UTRs) of three photosynthesis genes: *PsbS*, *Raf1*, and *SBPase*. We find that gene expression is most tunable within a ∼500 base pair core promoter region, where mutational effects are reproducible across biological replicates and predictive of protein output. Within these regions, we identify compact deletions and motif insertions that strongly increase protein production (>30-fold relative to wild type), exceeding the performance of transgenic enhancer elements. Mutation-effect relationships are gene-specific, highlighting the need for tailored regulatory maps. Our results establish a high-throughput strategy for *cis*-regulatory fine-mapping that enables crop improvements via minimal, precise, and non-transgenic gene edits.

## Introduction

The domestication and improvement of crops have historically relied on modifications to *cis*-regulatory DNA (crDNA), the portion of noncoding DNA involved in regulating gene expression^1^. While most flowering plants have retained a common set of genes and gene families^2,3^, modern plants have dramatically radiated by modifying the *cis*-regulatory components of promoters, enhancers, introns, and untranslated regions (UTRs)^4^. Advances in gene editing now enable precise modification of these *cis*-elements, a strategy termed quantitative trait engineering (QTE)^5^. Given the burgeoning regulatory acceptance of crop gene editing^6^, QTE has immense potential in tuning agronomically relevant phenotypes^7–9^. However, the promise of *cis*-regulatory engineering remains constrained by our limited understanding of plant regulatory architecture and low frequency of hypermorphic (activating) mutations.

This use of CRISPR-based QTE is particularly relevant to improving photosynthesis, a key target for crop productivity that suffers from suboptimal gene expression^10^. Although transgenic overexpression of key photosynthetic genes has enhanced yield and stress resilience in model systems^11–13^, regulatory and public acceptance barriers have precluded widespread field deployment. Non-transgenic upregulation via *cis*-regulatory editing could overcome this barrier, yet systematic identification of hypermorphic alleles in QTE experiments remains rare. For example, in a recent QTE screen targeting rice *PsbS*, only two hypermorphic alleles were discovered among 120 mutated promoters^14^.

To address this limitation, we developed a primary cell-based, massively parallel reporter assay (MPRA) in sorghum, a drought-resilient C4 bioenergy crop, to systematically map the effects of tens of thousands of CRISPR-mimicking mutations in the promoter and 5’ UTR sequences. Focusing on three key rate-limiting photosynthetic genes (*PsbS*, *SBPase*, *Raf1*), we quantified the effects of saturating deletions, substitutions, and motif insertions in their natural sequence context. Our results reveal promoter hotspots of regulatory control, identify compact sequence variants that drive strong transcriptional activation, and establish a generalizable strategy for regulatory tuning of photosynthetic genes.

## Results

### Systematically discovering gene expression-modulating promoter mutations using a protoplast massively parallel reporter assay

To understand how to improve photosynthetic gene expression using precise, small CRISPR-type edits (Fig. 1a), we developed an MPRA in sorghum protoplasts to identify hyper- and hypo-morphic mutations within *cis*-regulatory DNA (crDNA) (Fig. 1b). Briefly, tens of millions of healthy mesophyll protoplasts were isolated from partially etiolated *Sorghum bicolor cv. RTx430* seedlings (Extended Data Fig. 1, Methods), and transfected with plasmid libraries containing crDNA variants. After an 18 hour incubation in darkness, cells were exposed to 4 hours of light to stimulate photosynthetic gene expression. Next, barcoded mRNA was harvested, next-generation sequencing was used to quantify barcodes in the input DNA and output RNA, and linear regression was performed to infer individual variant effects, E_mut_ (Methods, Extended Data Fig. 2).

**Fig. 1.**
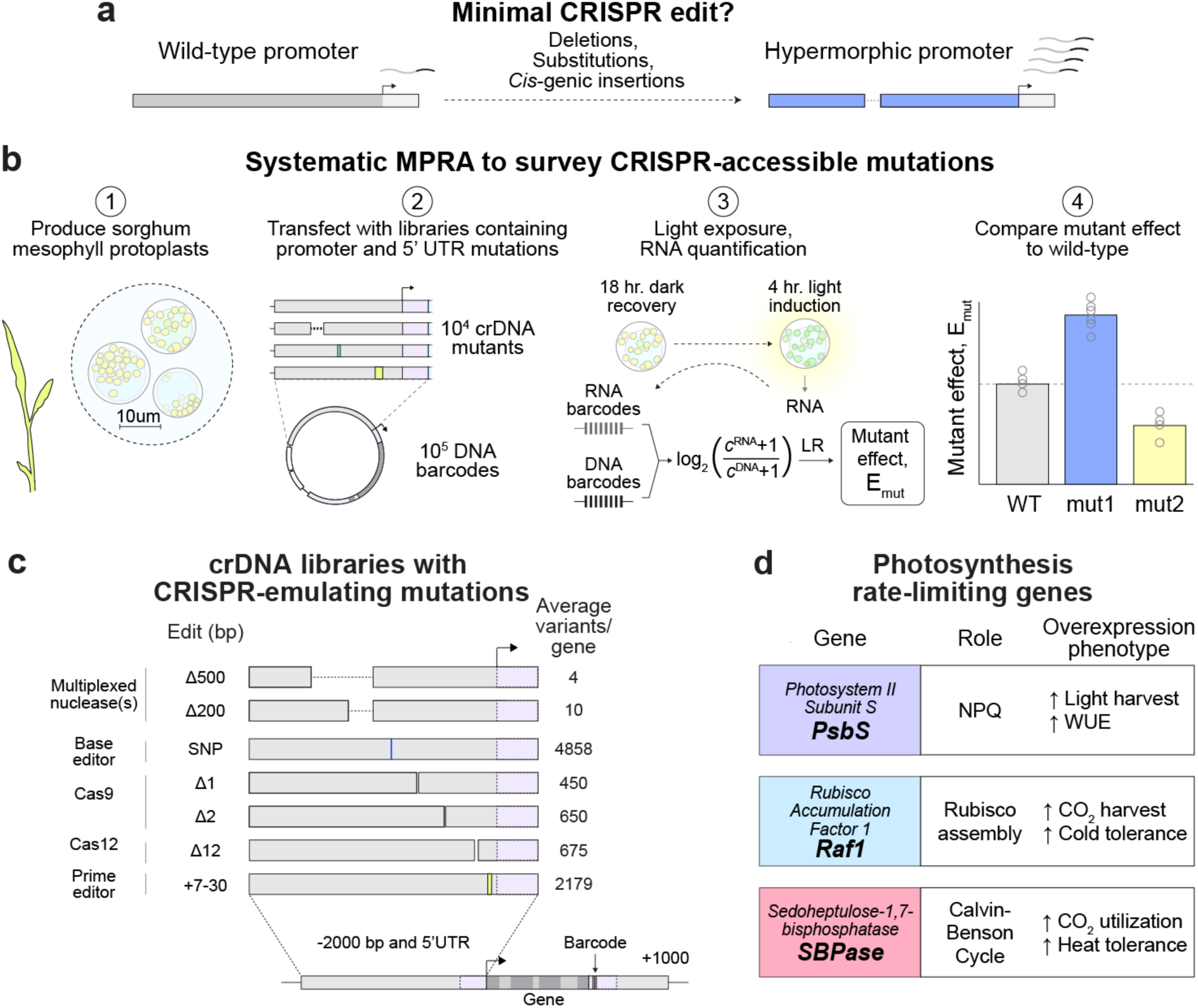
A massively parallel reporter assay for investigating CRISPR editing outcomes on sorghum *cis*-regulation. **a**, Depiction of MPRA aims, where saturated gene edits are surveyed at a given promoter/5’ UTR to discover the minimal edit required for gene expression tuning. **b**, Overview of MPRA steps. Protoplasts are produced from partially etiolated sorghum seedlings (1), then transfected with plasmid libraries containing 10^4^ cis-regulatory DNA (crDNA) mutants (2). After overnight incubation and light exposure, RNA is extracted from protoplast lysate and reverse-transcribed, then amplified alongside transfected DNA to measure mRNA expression (3). A linear regression is performed to infer individual mutant effects (E_mut_) on gene expression, and mutant effects (mut) are normalized relative to wild type expression (see Methods) (4). LR = linear regression. **c**, Sampled *cis*-regulatory DNA (crDNA) mutations include common editing outcomes of indicated CRISPR effectors. For each mutation type, its associated CRISPR effector, edit effect, and average number of variants/gene library is indicated. An average of ∼10,000 mutations per gene are designed and introduced into the 2 kb promoter region and 5’ UTR (variable length). 1 kb of downstream genomic sequence is included for each gene and a 15 bp DNA barcode is introduced 50 bp downstream of the stop codon. SNP = single nucleotide polymorphism. **d**, Surveyed genes account for major rate limits in photosynthetic carbon and light accumulation. Each gene - *PsbS*, *Raf1*, and *SBPase* - is listed alongside its photosynthetic role and its overexpression phenotype in transgenic studies^12,22,23,25–30^. NPQ = non-photochemical quenching, WUE = water use efficiency.

In contrast to prior work^15–17^, we focused our efforts on natural, 2 kilobase (kb) 5’ promoter as well as the 5’ untranslated region (5’ UTR) in the context of the full gene content, to account for possible dependencies of mutation effects on the intron and 3’ UTR sequences (Fig. 1c). Each crDNA library was expressed on a plasmid upstream of the relevant gene sequence with all introns followed by 1 kb of genomic sequence downstream of the stop codon. For each library, a 15 bp DNA barcode was introduced 50 bp downstream of the stop codon to allow for expression measurement. We also tested whether mutation effects can be discovered in a synthetic construct with the 2 kb promoter and 5’ UTR driving GFP expression with a synthetic terminator (Extended Data Fig. 3).

To mimic gene editing outcomes, we generated deletions (avg. 3398 deletions/library) that emulate CRISPR nuclease effects (Fig. 1c, Extended Data Fig. 3), including deletions produced by Type II/blunt endonucleases (avg. 1100 deletions/library), 12 bp deletions typical of Type V/staggered endonucleases (avg. 675 deletions/library), and non-overlapping 200 and 500 bp deletions simulating the effects of multiplex nuclease activity (∼14 deletions/library). Single nucleotide polymorphisms including A to G and T to C transitions within a 5 bp window were introduced to emulate base editor outcomes (avg. 4858 substitutions/library), and short insertions of *cis*-genic sorghum *cis*-regulatory DNA, which recapitulate motifs from expressed transcription factors^18^ and DNase I footprints^19^, were inserted to emulate sequence addition via prime editing or double-stranded oligodeoxynucleotide insertion^20^ (avg. 2179 insertions/library). Synthesis, amplification, and cloning of the oligopool library led to genetic variation beyond the designed library (e.g. deletions of unexpected size), which was characterized by long-read sequencing (see Methods).

To test our approach, we constructed libraries in three core photosynthesis genes expected to be rate-limiting in sorghum (Fig. 1d). *Photosystem II Subunit S (PsbS)* plays a central role in non-photochemical quenching^21^, and its transgenic overexpression in rice, soybean, and tobacco has been shown to improve light harvesting and agronomic yield^12,22,23^. *Rubisco accumulation factor 1 (Raf1)* is a molecular chaperone for the rubisco holoenzyme^24^ whose overexpression in wheat, maize, tomato, tobacco, and sorghum has been associated with improved carbon assimilation and biomass production^25–28^. Finally, *sedoheptulose-1,7,bisphosphatase (SBPase)* is a core enzyme in the Calvin-Benson cycle^10^ whose expression limits metabolic flux in regeneration of the rubisco substrate, ribulose 1,5-bisphosphate (RuBP)^29,30^. Our genes were also chosen for their diverse expression profiles both in terms of total and cell-specific expression (Supplementary Fig. 1). While *PsbS* is preferentially expressed in mesophyll cells, which make up the majority of leaf protoplasts^31,32^, *Raf1* and *SBPase* have both been co-opted for bundle-sheath specific expression in C4 sorghum. Despite these observed cell tropisms, all cells have some basal mesophyll expression (Supplementary Fig. 1).

### Systematic mutation profiling of the native *PsbS* promoter reveals reproducible gene expression tuning in the core promoter

We first applied our approach to study gene expression-modulating mutations to the natural crDNA of mesophyll-expressed *PsbS*. The gene expression effects of 10,096 mutations across the 2 kb promoter and 5’ UTR reveal a ‘core’ promoter region (−400 bp to the start codon) of increased gene expression variance, compared to a ‘distal’ promoter region of decreased variance (<-400 bp) (Fig. 2a). This is consistent with reported accessibility measurements, where accessibility diminishes ∼400 bp upstream of expressed genes^33,34^. The reproducibility between biological replicates is high for mutations in this core promoter (Pearson r: 0.68), in contrast to the distal promoter (>-400 bp, Pearson r: 0.30) (Fig. 2b-c). Deletions within the core promoter reveal slightly higher replicate correlations (Pearson r: 0.75) (Fig. 2d), and insertions within the core region have slightly lower correlation (Pearson r: 0.62) (Fig. 2e).

**Fig. 2.**
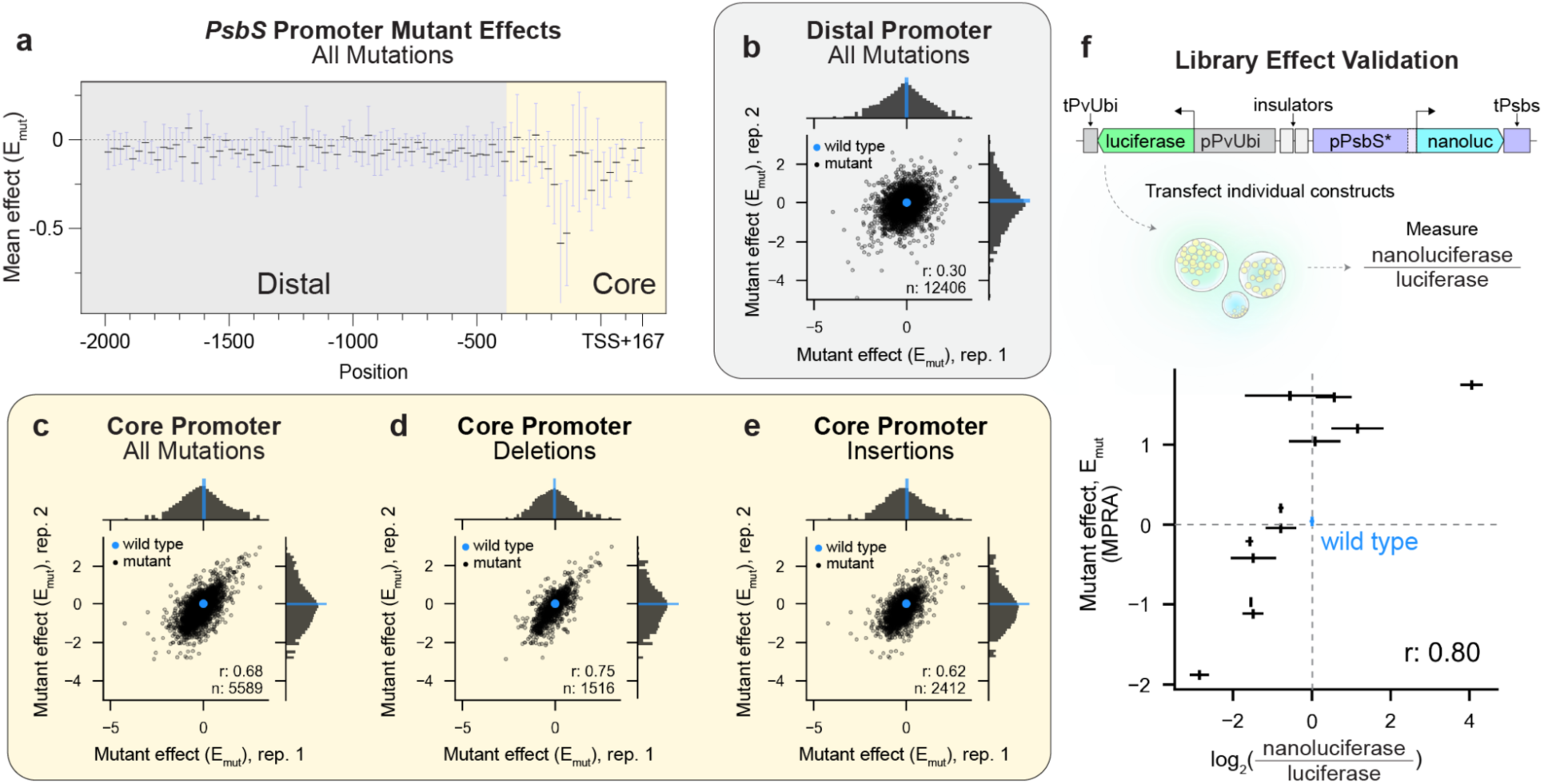
Systematic mutagenesis of *PsbS cis*-regulatory DNA reveals reproducible gene expression tuning in the core promoter. **a**, The mean and variance of all mutation effects, E_mut_, across the *PsbS cis*-regulatory DNA is shown indicating the core promoter region (−400 to +168 bp from the transcriptional start site; E_mut_ in log_2_ scale). **b-e**, The biological replicate reproducibility is shown for (**b**) the ‘distal’ promoter (defined as <-400 bp from TSS), (**c**) all mutations, (**d**) deletions, and (**e**) insertions within the ‘core’ promoter (defined as >-400 from TSS) and 5’ UTR. Pearson correlation coefficients (r) and number of mutations (n) for each set are indicated. **f**, 12 promoter and 5’ UTR mutants (pPsbS*) identified in the MPRA are validated for their effects in expressing nanoluciferase (cyan) relative to constitutive firefly luciferase (green) expressed with a switchgrass ubiquitin promoter (pPvUbi) and terminator (tPvUbi). The expression cassettes are separated with an *Arabidopsis* AT-5-IV-2 insulator and a maize IN2-1 terminator. For nanoluciferase/luciferase measurements n = 4 technical replicates from 2-3 independent seed batches. The Pearson correlation coefficient, r, is indicated.

To benchmark our system, we tested whether these library-measured transcriptional effects correlate with changes in protein production. To do so, we picked 12 deletions and insertions that span the range of library mutant effects and tested their ability to drive expression of nanoluciferase in an orthogonal enzymatic assay^35^. This bulk assay works by measuring nanoluciferase production from a mutant promoter compared to a constitutively driven firefly luciferase expression from the same plasmid in order to normalize for transformation and protein production efficiency. We found a high correlation between our library effect measurements and the nanoluciferase protein production levels in sampled promoter/5’ UTR mutants (Pearson r: 0.80, Fig. 2f). The protein production correlation of these same mutation effects observed in our synthetic library driving GFP expression was low (Pearson r: −0.26, Extended Data Fig. 4g), demonstrating a context dependence between mutation and endogenous gene structure. Together, these results indicate that systematic library measurements of CRISPR-mimicking promoter mutation effects are meaningful and reproducible for perturbations in the core promoter, and that these effects translate to protein production.

### Mutational scanning of the *PsbS* promoter reveal hotspots for expression regulation

We next investigated which elements of the *PsbS* core promoter are responsible for regulating expression. To probe this, we first inspected our genetic deletions for their effects on *PsbS* transcription. Here, we observed that hypomorphic deletions cluster around the region −180 to −120, indicating its core function in *PsbS* transcription (Fig. 3a). We note that a 12 bp deletion at position −168 is more hypomorphic than the removal of an entire −201 bp region at position −200 (Fig. 3b).

**Fig. 3.**
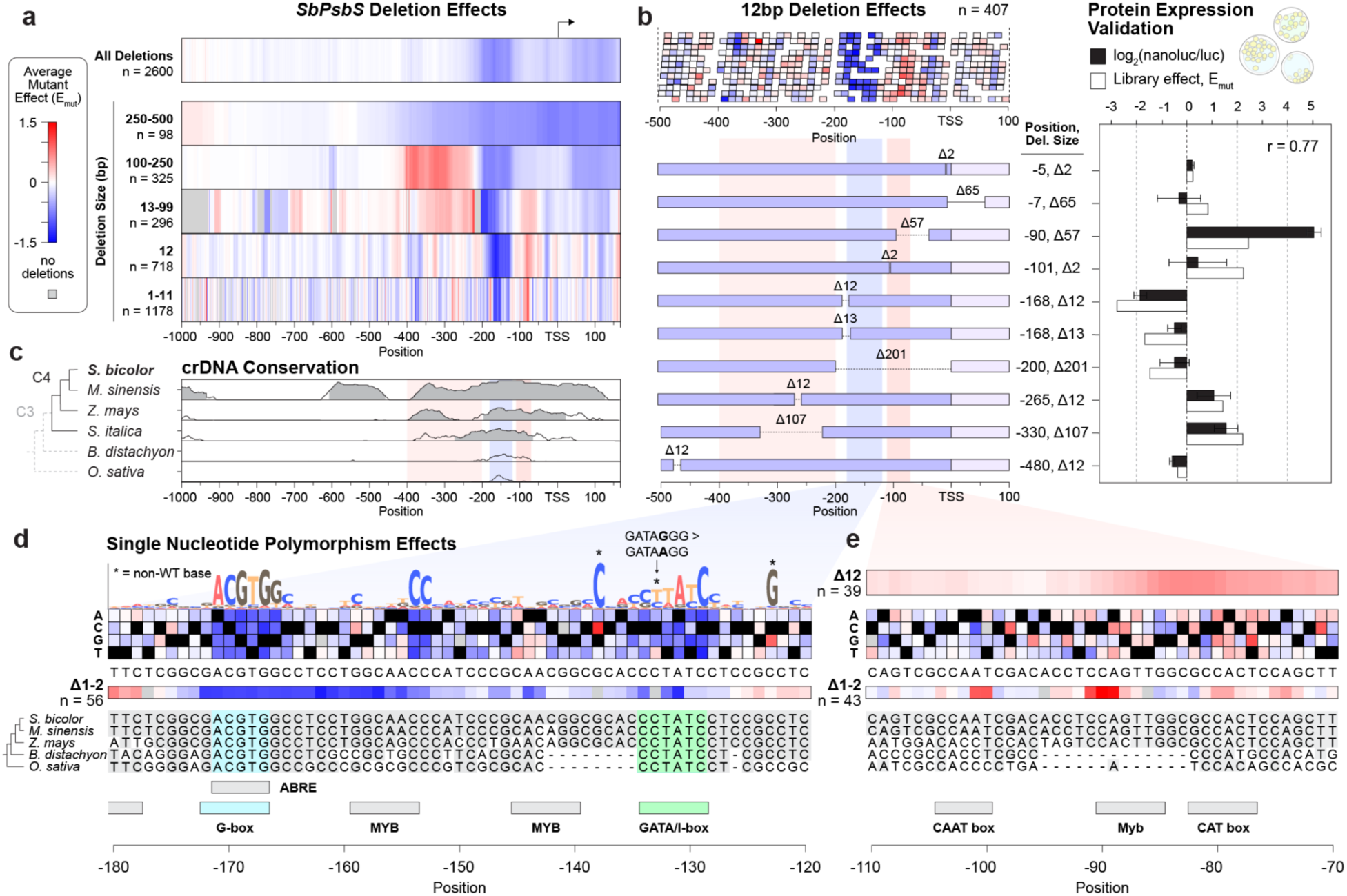
Saturated mutation profiling of *SbPsbS* crDNA reveals how to tune gene expression with deletions and substitutions. **a**, The average effect of deletions of various lengths overlapping a given position across the *PsbS* promoter from −1000 bp from the TSS through the full 5’ UTR. Deletions are grouped by their size as indicated on the y-axis with the number of deletions (n) in each group displayed. At each position, the average deletion effects (E_mut_) is displayed according to the color bar. **b**, The effect of individual 12 bp deletions in the region −500 to 100 is shown (top). Below, validated deletions in the region −500 to 100 bp are displayed in terms of their library measured effect, E_mut_, (white bars) and normalized log_2_(nanoluciferase/luciferase) effect (black bars, 2 biological replicates with 2 technical replicates each). The Pearson correlation coefficient, r, between library and nanoluciferase measurements is indicated. **c,** Smoothed conservation tracks (100 bp sliding window) for the *SbPsbS* promoter compared to 5 monocot orthologs are shown. Grey shaded regions indicate sites of high sequence similarity (>70% for at least 100 bp). Identified regions of hypermorphic (red) and hypomorphic (blue) deletion effects are shaded. Y-axis ranges from 50 to 100% conservation in all tracks. **d,** Individual substitution effects and 1-2 bp average deletion effects in the promoter region −180 to −120 are shown. Species-level alignments from the *PsbS* promoter are displayed with conserved bases highlighted in grey. Sequence logos of base preferences are shown (Methods) with deviations from wild type bases indicated with asterisks (top). An activating substitution at position −133 is highlighted, showing an activating reversion to a canonical I-box motif (GATAAGG)^38^. Predicted TFBSs are displayed (see Methods). **e,** 12 bp deletion average mutant effect, substitution effects, and small deletion effects are shown in the region −110 to −70 overlapping top hypermorphic deletions. Heatmap color scale for all subfigures is shown in panel **a**.

This necessary promoter region is flanked by two stretches enriched for hypermorphic deletions. These hypermorphic regions differ in their deletion size response: on average, large deletions (100-250 bp) in the region −400 to −200 from the TSS activate transcription, but smaller deletions (<100 bp) in the same area have a modest if not negligible effect on expression (Fig. 3a). Conversely, small (<100 bp) deletions lead to overexpression in the −110 to −70 region (Fig. 3a), presumably constrained by core sites of transcription initiation, with the strongest hypermorphic deletion being a 57 bp deletion starting 90 bp upstream of the transcription start site (Fig. 3a-b). To corroborate our findings, we performed orthogonal protein production measurements for these promoter and UTR deletions (Fig. 3b) and found that deletion effects correlated with protein production (Pearson r: 0.77).

Sequence conservation and chromatin accessibility are common heuristics for designing and explaining promoter mutation experiments, so we wondered whether these metrics could predict our mutational scanning results. We find that our most repressive hypomorphic deletions correspond with a highly conserved noncoding region across related grass species predating the transition to C4 (Fig. 3c, blue box), and overlap with the local maximum of a large (∼600 bp) mesophyll and bundle sheath accessible chromatin region (Supplementary Fig. 1b-d)^36^.

To determine the dependence of *PsbS* expression on particular motifs at higher resolution, we examined the effects of 4505 substitutions and 1111 1-2 bp deletions. Many single nucleotide variants and small deletions that cause loss of expression of *PsbS* are found at −169 and −134 (Fig. 3d). Comparing these results to predicted transcription factor binding sites (TFBS)^37^ revealed that these sites directly overlap with highly conserved, reverse-strand G-box and I-box motifs. These have been demonstrated to be essential for the light-mediated activation of photosynthesis genes in C4 plants^38^. Although both motifs occur multiple times throughout the *PsbS* upstream region, no other instances of these motifs affect *PsbS* expression when mutated (Extended Data Fig. 5). In certain positions, substitutions that modified putative core motifs were slightly more activating than wild type sequences (Fig. 3d, indicated by asterisks), such as a C>T transition that modified the native I-box-like sequence (reverse GATAGGG at position - 133) to its more canonical form in light-responsive genes (reverse GATAAGG)^38,39^.

We carried out a similar inspection of the region containing the most hypermorphic deletions (−110 to −70) (Fig. 3b red boxes, Fig. 3e). Strong-effect deletions (>12 bp) in the region overlapping a C4-specific binding site for a Myb/bZIP TF (CAGTTG) and a CAT box (GCCACT) were particularly activating, though no single substitution or indel mutation (Fig. 3e) was as hypermorphic as larger deletions (e.g. the top hypermorphic 57 bp deletion at position −90) in this region (Fig. 3b), perhaps indicating the involvement of multiple co-localized repressive elements or the activating potential of rearranging core enhancer elements closer to the TSS.

As rice and sorghum contain a conserved region for transcriptional activation, we investigated the extent to which rice protoplasts could recapitulate observed differences in sorghum *PsbS* promoter activity. We tested our *SbPsbS* nanoluciferase variants in light-grown rice protoplasts and found that protein production of key *PsbS* mutations was correlated with sorghum MPRA values (Pearson r: 0.59) and sorghum protein production (Pearson r: 0.89) (Extended Data Fig. 6d). Approximately an order or magnitude more protoplasts can be obtained from light-grown rice than etiolated sorghum, so we used our rice assay to study how nanoluciferase production responded over a 4-hour light induction period following a 12-hour dark incubation (Extended Data Fig. 6e). Relative protein production between constructs was constant regardless of light induction, though all mutant constructs increased expression throughout the time course.

A previous promoter mutagenesis experiment in rice allowed us to compare our mutants, particularly hypomorphs, to *in planta* results^14^. The sorghum hypomorphic gene expression hotspot (−180 to −120) is highly conserved in rice (Fig. 3b), and when mutagenized has a corresponding hypomorphic NPQ effect in mature plants (Extended Data Fig. 6a-c). In rice, deletions in the G-box induce a decreased NPQ phenotype, while deletion of both the G- and I-box phenocopies a *PsbS* knockout. This suggests that our library measurements correspond to *in planta* phenotypes and that the surveyed set of hypomorphic alleles can translate to the full range of *PsbS* downregulation.

### Mutations that mimic prime editing outcomes reveal hypermorphic motif insertions and their positional dependence

We next wondered whether prime edit-like insertions of small motifs can enable overexpression, and how such perturbations compare to deletions as a tool for modulating gene expression. To do so, we examined the effect of 80 different motifs (length 8 to 25 bp) inserted every 5 bp from −150 to +45 relative to the transcriptional start site, in both forward and reverse orientation (Fig. 4a). As described in the Methods, these were selected based on imputed binding motifs of sorghum leaf-expressed transcription factors^40^, as well as DNase I footprints^19^.

**Fig. 4.**
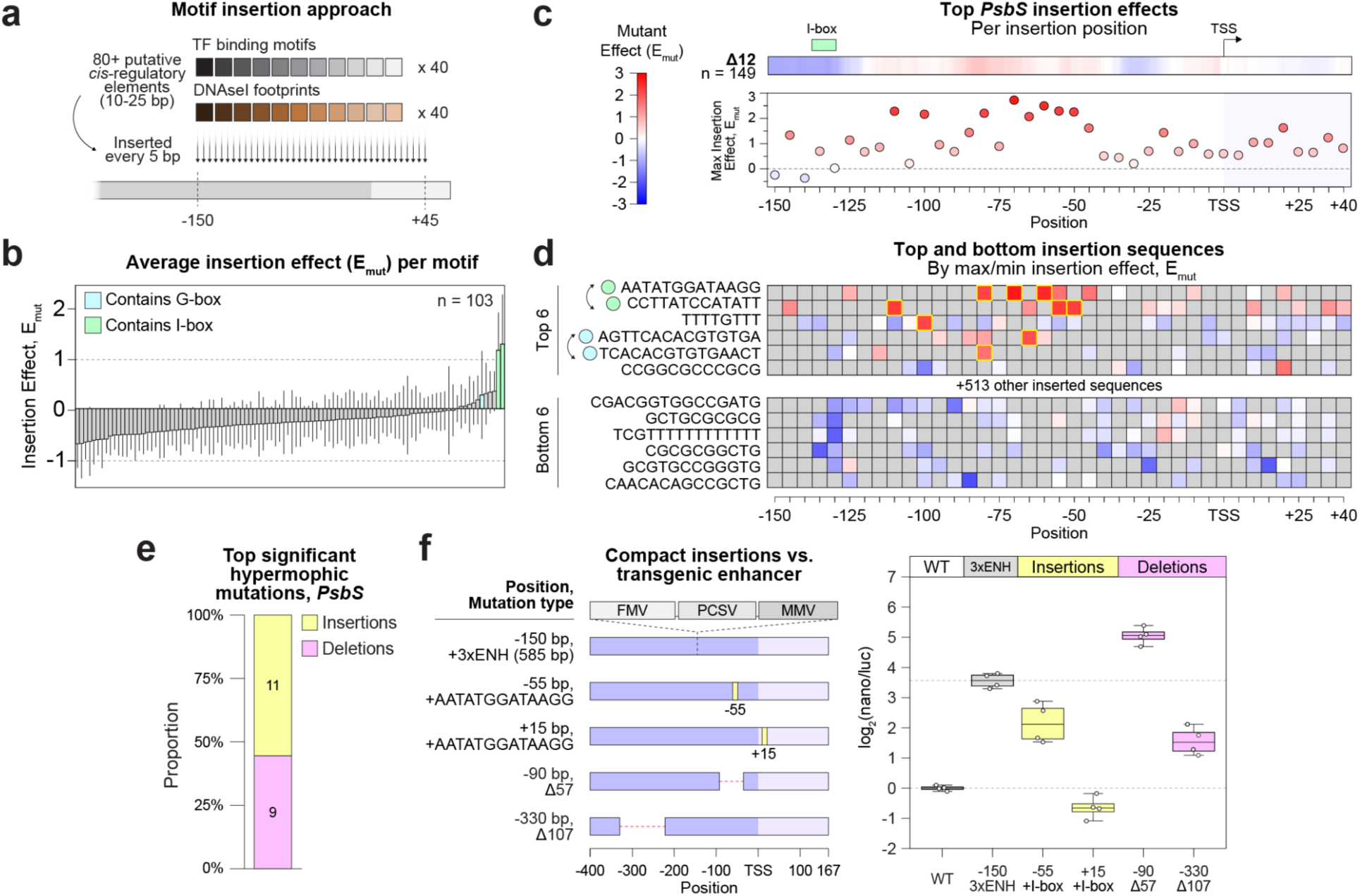
Systematic insertions in *SbPsbS* crDNA reveals compact motifs for expression enhancement. **a**, Diagram of motif screening by inserting motifs every 5 bp between the −150 and +45 region relative to the TSS. **b**, Average effect of 103 insertion sequences with >7 measurements in the *PsbS* promoter. Motifs that contain G-boxes (CACGTG) or I-boxes (GATAAGG) are indicated by cyan or green shading, respectively. **c**, Maximum enhancer effect by position is plotted for each 5 bp position between −150 and +45. The average effect of 12 bp deletions is shown above. **d**, The top and bottom 6 insertions by maximum and minimum effect are displayed at all sampled positions, with arrows indicating reverse complements. Motifs that contain G-box (cyan) or I-box (green) sequences are indicated, and individual significant and meaningful insertions (Bonferroni corrected p-value < 0.05, mutation effect, E_mut_ > 1.5, and mean log read ratio of reads containing this mutation > 1.5, Methods) are highlighted in yellow. **e**, Fraction of top significant and meaningful mutations in the *PsbS* promoter by type (deletions, insertions, and substitutions) are shown. **f**, The effect of top deletions and insertions in the *PsbS* promoter is compared to a transgenic enhancer, 3xENH (585 bp) using nanoluciferase protein production measurements (n = 4). Heatmap color scale for subfigures **c** and **d** is shown in panel **c**.

We observed three unique motifs across 11 specific insertions, which cause significant and meaningful overexpression of *PsbS* (Bonferroni corrected p-value < 0.05, effect size E_mut_ > 1.5, mean log read ratio of raw reads > 1.5, see Methods, Fig. 4b), though the majority of insertions have no or deleterious expression effects. The specific effects across the promoter reveal that expression-increasing motif insertions (AATATGGATAAGG or its reverse complement) containing the I-box (GATAAGG) work nonspecifically when inserted in the region −110 to −45 (Fig. 4d, top 2 rows), overlapping our strongest characterized hypermorphic deletions. Inserting such I-box motifs outside of this region does not necessarily lead to increases in expression (Fig. 4c-d). Similarly, G-box containing motifs were also activating in this region in forward and reverse orientation (Fig. 4d, rows 4 and 5). In contrast, the motif insertion of TTTTGTTT only increases expression of *PsbS* when inserted at position −100 (Fig. 4d, row 3).

We next investigated how these motif insertions compare to deletions for achieving *PsbS* overexpression, as well as to a 585 bp industry-standard enhancer (3xENH) used for sorghum overexpression, which is composed of three viral enhancer sequences^41^. The significantly hypermorphic perturbations in *PsbS* include 9 deletions and 11 motif insertions (Fig. 4e). Interestingly, the top deletion (Δ57 bp at position −90 bp from the TSS: 5.05 log_2_ fold increase, ∼33 fold increase) outperforms the top I-box motif insertion at position −55 bp (2.16 log2 fold increase, ∼4.5 fold increase) as well as the 3xENH sequence (3.56 log_2_ fold increase, ∼12 fold increase) in achieving protein overexpression (Fig. 4f). These results indicate that our tested motif insertions can cause overexpression in a position-dependent manner, but only a deletion can outperform industry-standard, transgenic approaches in causing overexpression of *PsbS*.

### Systematic mutation profiling reveals gene-specific expression modulation

To assess whether our specific findings about *PsbS* expression modulation hold in other promoters, we also applied a systematic mutation profiling approach to *Raf1* and *SBPase* (7,796 and 13,974 mutations in the promoter and 5’ UTR of *Raf1* and *SBPase*, respectively). Similarly to *PsbS*, we found that a narrow, core promoter region (−400 onwards) around the transcriptional start site shows increased mutation effect variance (Fig. 5a) and that the replicate reproducibility is higher in this region compared to the distal promoter from −2000 to −400 bp from the transcriptional start site (Extended Data 4).

**Fig. 5.**
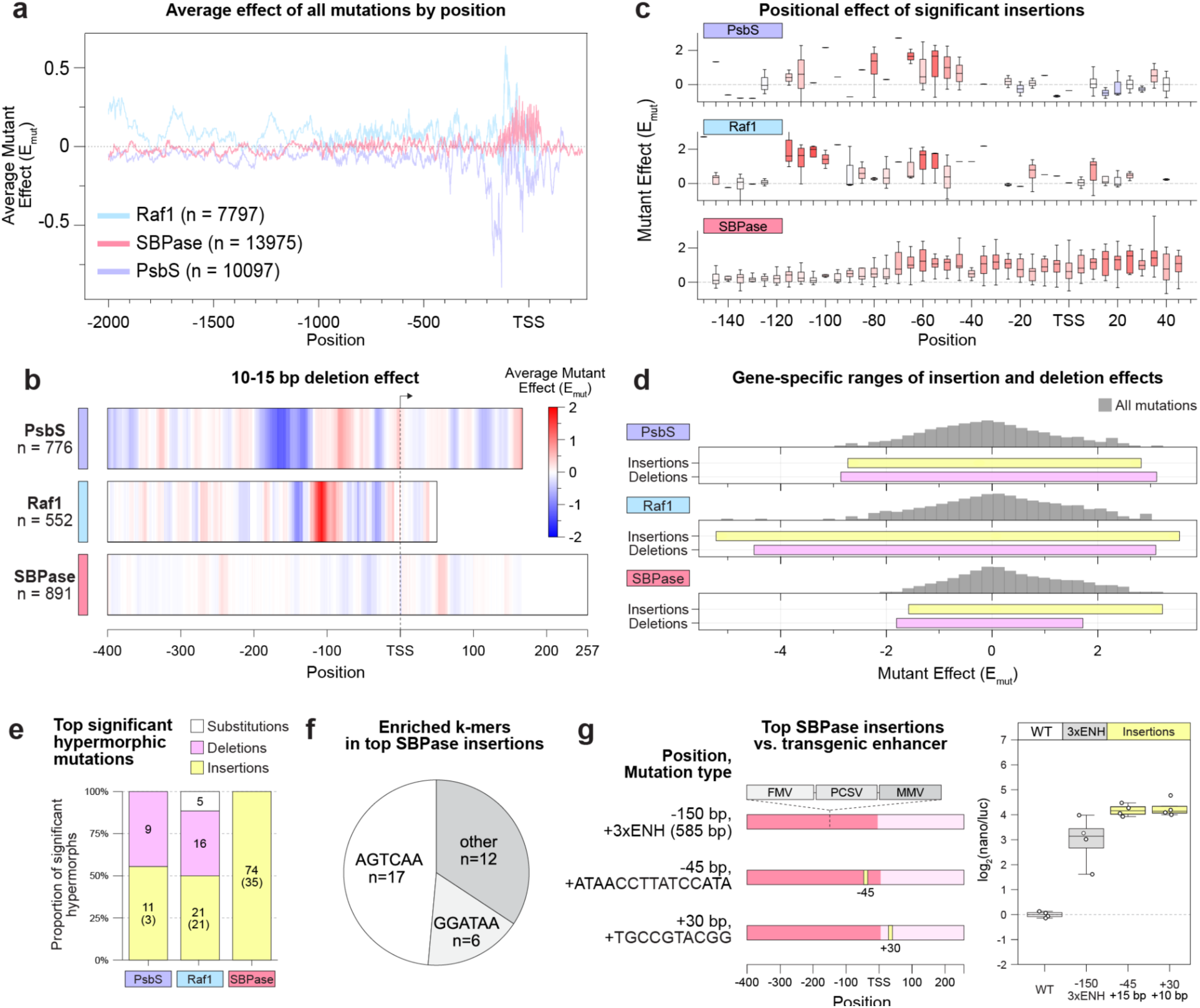
Mutation scanning reveals gene-specific expression modulation patterns. **a**, The rolling average effect of all mutations for three genes is displayed, identifying a core promoter region (−400 onwards) with marked gene expression effects across all genes. **b**, The average effect of 10-15 bp deletions is displayed across the core promoter and 5’ UTR for three genes. **c**, The effect of significant and meaningful hypermorphic insertions (Bonferroni corrected p-value < 0.05, mutation effect, E_mut_ > 1.5, and mean log read ratio of reads containing this mutation > 1.5, Methods) across positions are shown for three genes. For each position, the boxplot of mutation effects, E_mut_, with median lines, boxes representing interquartile ranges (IQR), and whiskers representing the range up to 1.5 of the IQR shown. **d**, The range of effects for insertions or deletions is shown for each gene underneath the log-density of all mutation effects. **e**, The mutation types of significant and meaningful hypermorphic mutations are displayed for three genes. The number of instances for each mutation type is indicated, with the number of unique motifs indicated in parentheses. **f,** The top enriched submotifs among unique *SBPase* overexpressing motifs are displayed. **g**, Protein production levels due to *SBPase* overexpressing insertions are validated and compared against a transgenic 3xENH enhancer insertion using nanoluciferase/luciferase measurements (n = 4).

Examining specific effects of 10-15 bp deletions between these core promoters revealed distinct differences in *cis*-regulatory structure. For example, lowly-expressed *Raf1* can be upregulated via deletions in a small window (−120 to −90) that is distinct from the hypermorphic deletion windows in *PsbS* (−400 to 200 and −110 to −50, Fig. 5b, middle). In contrast, the *SBPase* library contains no significant hypo- or hyper-morphic deletions (Fig. 5b, bottom, Supplementary Fig. 2).

Motif insertions also show differences in their position-dependent effects across genes. Examining the distribution of significant insertions for each gene reveals that *SBPase* can be overexpressed with motif insertions within the −70 to +45 window, whereas motif insertions in *Raf1* and *PsbS* are only activating within the −120 to −30 region (Fig. 5c).

Across these genes, the type of mutation capable of inducing significant overexpression differed (Fig. 5d-e). While significant upregulation of *PsbS* and *Raf1* can be accomplished with both insertions and deletions, *SBPase* upregulation could only be observed with motif insertions but not deletions (Fig. 5d). In *SBPase*, we uncovered 35 unique motifs across 74 insertion instances that cause significant and meaningful overexpression (Fig. 5e). Around two thirds (23/35) of these 35 unique motifs contain either an AGTCAA or GGATAA (I-box-like) submotif (Fig. 5f). For *SBPase*, we also compared our top overexpressing motif insertions against the 3xENH enhancer sequence. A 10-bp insertion at +30 and a 15-bp dual-I-box insertion at −45 both overexpressed (>4 log_2_-fold, >16-fold), beyond the 585 bp 3xENH enhancer insertion (∼3 log_2_-fold, ∼8-fold) (Fig. 5g).

Together, these results suggest that our approach can reproducibly reveal the promoter-specific mutation effects for modulating gene expression across different genes, and that, while core-promoter dependency is shared among these three promoters, the details of how to accomplish expression modulation vary markedly between them.

### A fine-tuned genomic language model predicts core promoter deletion effects for PsbS

We wondered to what extent the observed mutational outcomes could be predicted from sequence context without having to perform such extensive variant effect measurements. While current machine learning models trained on promoter sequences and RNA-seq data, without variant-level MPRA data, can robustly predict expression variation across genes, prediction of higher resolution variant effects for a given gene remains challenging^42–44^. We hypothesized that a moderate level of predictive performance might still be achievable, which could motivate further investigation into model training and data collection strategies.

We fine-tuned the genomic language model GPN^45^ to predict RNA-seq expression levels from promoter sequences (−256 to +256 bp from the transcriptional start site) across sorghum leaf tissue (Fig. 6a). Model performance across transcribed genes was not indicative of performance across MPRA variants (Extended Data Fig. 8a). Consequently, we validated such fine-tuned models on measured single-nucleotide variants, and tested model performance on the remaining variant measurements.

**Fig. 6.**
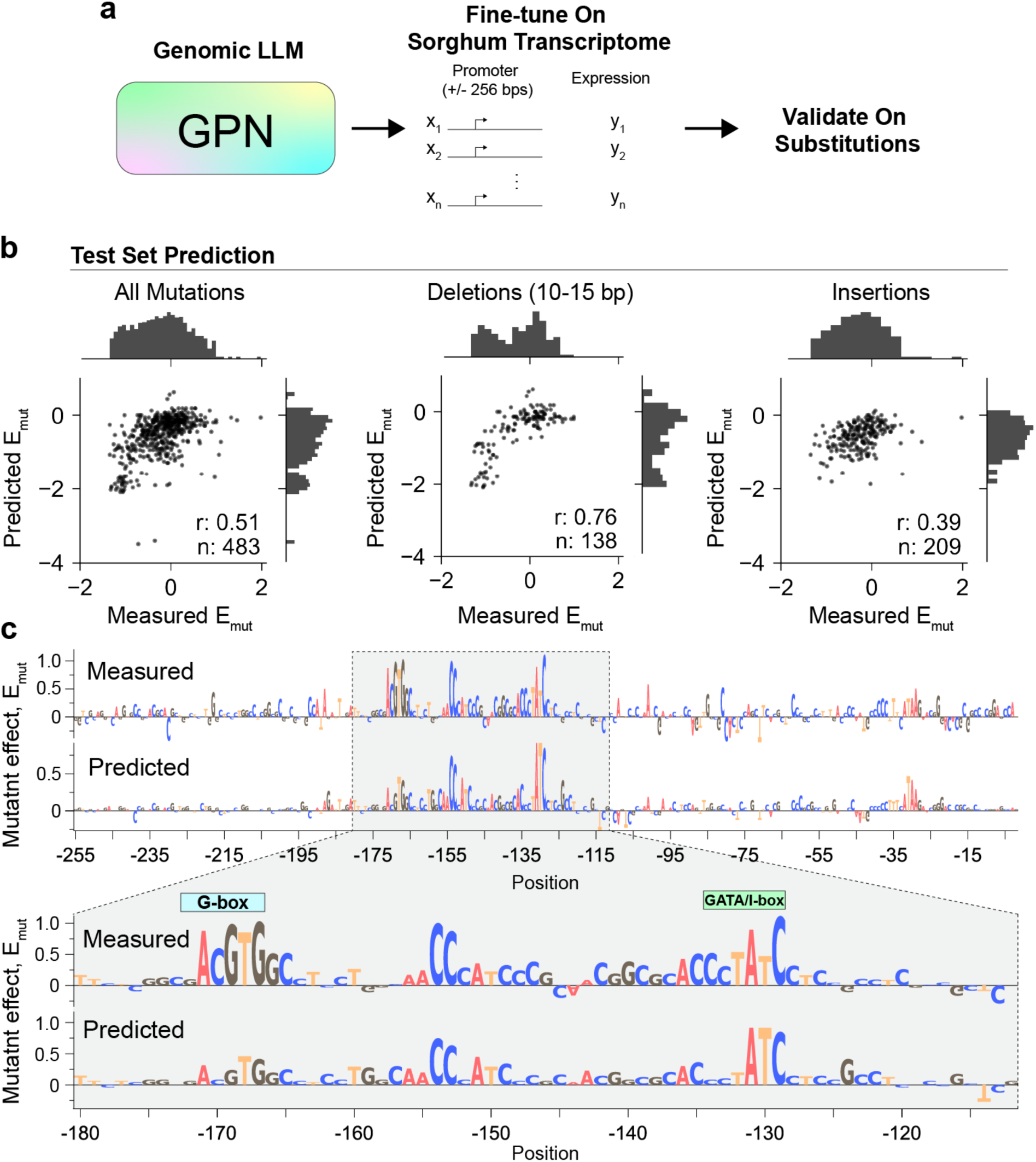
A fine-tuned genomic language model predicts core promoter deletion effects for *PsbS.* **a**, GPN was fine-tuned on sorghum transcriptome data (using +/- 256 bp of promoter sequence) and model validation was performed using library-measured single nucleotide substitution effects. **b**, Test set predictions from the model are shown for all mutations (left), deletions (middle), and insertions (right). The Pearson correlation coefficient, r, and number of observations, n, are indicated. **c**, Logo plots for the measured single nucleotide substitution effects vs. the model predicted substitution effects from *in silico* mutagenesis are shown.

We found a modest correlation between predicted and observed test set mutation effects across the −256 to TSS region in *PsbS* (Pearson r: 0.51, Fig. 6b left). We detect increased correlation for medium sized (10-15 bp) deletions that are disruptive to expression (Pearson r: 0.76, Fig. 6b middle), with less correlation for motif insertions (Pearson r: 0.39, Fig. 6b right) or activating variants. Furthermore, *in silico* mutagenesis reveals effects similar to measured single-nucleotide variants (Fig. 6c). We note that variant effect prediction results for *Raf1* and *SBPase* genes were significantly weaker (Extended Data Fig. 8b). We hypothesize that this results from the low expression of these genes in our assayed tissue, coupled with our observation that model predictions are best at predicting decreased gene expression. Together, these results suggest that genomic language models have the potential to identify essential regulatory motifs that maintain gene expression, but our MPRA approach is particularly capable of identifying hypermorphic crDNA mutations.

## Discussion

Gene editing holds enormous promise for precise, rapid, and regulation-compliant crop improvement. However, its application to gene expression tuning is constrained by an incomplete understanding of the functional architecture of plant promoters. Existing QTE strategies often rely on heuristics such as sequence conservation or chromatin accessibility, which incompletely capture regulatory logic^46,47^, lack basepair resolution, and give no prediction on the directionality of mutation effects. Here, we provide a comprehensive map of expression-modulating *cis*-regulatory variants across native promoter and 5’ UTR regions to systematically determine the effect of CRISPR-type editing outcomes. While experimental approaches to mutagenize promoters/UTRs *in planta* infrequently find hypermorphic variation^14^, this systematic approach enables identification of hundreds of expression-modulating mutations. We find that regulatory variants are strongest within a ∼500 bp “core promoter” window surrounding the transcriptional start site, where mutational effects are reproducible and predictive of protein expression. Although these data are focused on only a small number of genes, the observed mutation-expression relationships are apparently gene-specific. These findings underscore the need for gene-specific *cis*-regulatory maps to guide quantitative trait engineering (QTE).

In particular, this approach enables testing allelic variation beyond what exists in extant breeding populations. This can be especially useful in sorghum, which is speculated to have limited nucleotide diversity in noncoding DNA compared to maize and other grasses^48^. For example, while GWAS studies to identify the genetic drivers of NPQ efficiency in *Arabidopsis*^49^, rice^50^, and maize^51^ have all identified noncoding *PsbS* variants, such efforts have failed to identify *SbPsbS* in GWAS studies on sorghum NPQ^52^ and photosynthetic efficiency^53^. These observations suggest either that *SbPsbS* alleles for photosynthetic enhancement are rare or nonexistent in standing genetic variation, or that *SbPsbS* expression levels cannot be altered to improve photosynthetic efficiency. Studies demonstrating that transgenic overexpression of *PsbS* improves photosynthesis suggest the former^12,14^.

Despite numerous studies in humans and plants demonstrating reproducibility between cell-based MPRAs and organism-level data^15,35,54,55^, the limits of which types of promoters and their spatiotemporal characteristics can be faithfully studied in protoplast assays remains to be elucidated. As such, we have focused on photosynthetic genes, whose desired overexpression is constitutive and non-tissue specific, and whose regulatory logic has historically been captured via protoplast assay^56,57^. We note that our top hypomorphic deletions in the *PsbS* promoter show corresponding phenotypic effects in rice (Extended Data Fig. 6), but future work will be needed to validate our hypermorphic mutations *in planta*.

Unlike prior efforts limited to small synthetic constructs^16,58,59^, our approach enables fine-scale dissection of native regulatory regions, revealing gain- and loss-of-function variants across three endogenous sorghum promoters to identify promising candidates for tuning native gene expression via CRISPR. These variants span diverse mutation classes, including short deletions, base substitutions, and motif insertions, and recapitulate known functional motifs such as G- and I-boxes. Importantly, our top activating deletions and insertions outperform transgenic enhancers - at least at the tested position - in driving expression, highlighting their potential for non-transgenic crop improvement.

Our results can also be used to understand the evolution of regulatory modules. For example, the most hypomorphic mutations in sorghum *PsbS* occur at a deeply conserved promoter region (−172 to −125 bp), suggesting its conserved function in monocot *PsbS* expression pre-dating the transition to C4. Conversely, the strongest activating deletions in *PsbS* and *Raf1* cluster in C4-specific promoter regions (Fig. 3b, Supplementary Fig. 2), perhaps indicating their role in cell-type specific *cis*-regulatory rewiring during C4 photosynthetic evolution. Whether mutations like our strongest *PsbS* deletion (57 bp at position −90 from the TSS) disrupt C4/C3-specific repressive modules, reconfigure core enhancer regions, or create *de novo* activating elements is an area for future mechanistic investigation.

We observe potential for genomic language models in conjunction with transcriptome data to predict our observed variant effects in the *PsbS* promoter without reliance on extensive MPRA data, but note that our current approach lacks robustness across genes and even random initializations. As one could expect, these models trained on natural promoters fail to identify hypermorphic mutation effects, which arise from significantly sized deletions or motif insertions, emphasizing the utility of MPRA measurements to determine mutation effects beyond variation found in natural sequences. We suspect that significant improvements to variant effect predictions could come from exploration of training strategies that leverage both transcriptome and increased variant effect measurements across promoter architectures, extending prior efforts in learning predictive functions of promoter outputs from MPRAs^58^.

In sum, we show that this high-throughput framework can be extended across genes, mutation types, and expression outcomes. It provides a scalable platform for identifying minimal, non-transgenic edits that modulate gene expression, and can inform rational design of editing strategies in crops. By bridging *cis*-regulatory genomics and functional variant screening, this approach stands to enable a new era of precise, multiplexed engineering of complex traits.

## Methods

### Sorghum cultivation and protoplast generation

*Sorghum bicolor cv. RTx430* isolation was adapted from maize protoplast protocols^55,60,61^ and optimized for application in etiolated sorghum seedlings. Seeds were sterilized (70% EtOH 5 minutes, 30% NaOCl + 0.1% Tween-20 20 minutes, three washes with ddH_2_O 5 minutes each) and germinated for 48 hours on damp filter paper in the dark at 28°C. Generally, larger seeds are more suitable for protoplasting. Once germinated, seedlings with coleoptiles 2-3 cm were gently transplanted into 36 mm peat plugs and grown in the light (100 μmol m^−2^ s^−1^, 16:8 photoperiod, 28°C, 80% relative humidity) for 2-3 days to inhibit mesocotyl elongation. As soon as plants began developing a third leaf (i.e. the second leaf after the coleoptile leaf), or once they began noticeably greening, they were moved to a darkened growth chamber (28°C, 60% relative humidity) to develop for 8-10 days. All plants must be maximally etiolated for viable protoplast harvest, so any seedlings that were not fully yellow at the time of protoplasting were discarded, as were any that displayed any foliar abnormalities.

Plants were harvested when the third leaf was maximally extended, when plant height was 15-25 cm. Turgid, healthy third leaves were detached from plants with a razor blade and their base and tip were removed, leaving a ∼10 cm section. Approximately ten sections at a time were bundled and dissected into 0.5 mm strips perpendicular to the vein on clean paper. Any leaf strips that produced excess moisture on the paper when sliced were discarded. Once cut, strips were immediately transferred to 50-75 mL aliquots of fresh digestive solution (10 mM KCl, 0.5 M mannitol, 8 mM MES pH 5.7, 1 mM CaCl_2_, 1.5% w/v cellulase, 0.75% w/v macerozyme, 0.1% w/v BSA) in 200 mL beakers covered in foil to fully exclude light (∼25-35 leaves/beaker). Digesting strips were vacuum infiltrated (15 inHg, RT) for 3-5 minutes and incubated in a dark shaking incubator (40 rpm, 28°C) for 6 hours, then mixed with an an equal volume of W5 solution (154 mM NaCl, 125 mM CaCl_2_, 5 mM KCl, 2 mM MES pH 5.7) and incubated (80 rpm, 28°C) for 1 hour. Digestive solution was filtered with 40 μm cell strainers into 50 mL conical tubes, then spun down at 100 x g for 5 minutes. Yellow cell pellets were consolidated into 1-2 50 mL conical tubes and resuspended in 5-10 mL W5 solution, then checked for viability using Evan’s Blue dye. Any preparation that had more than 15% stained cells was discarded. Cells were spun down under the same conditions and resuspended in fresh MMG solution (0.4 M mannitol, 15 mM MgCl_2_, 4 mM MES pH 5.7) at a concentration of 2.5 x 10^6^ cells/mL.

To transfect libraries, 1.2 mL of cells (3 x 10^6^ cells total) in 15 mL conical tubes were combined with 100 μg purified plasmid in 200 μL of H_2_O and an equal volume (1.4 mL) of fresh, filtered PEG-CaCl_2_ solution (0.2 M mannitol, 0.1 M CaCl_2_, 40% w/v PEG-4000). Tubes were mixed by gently rocking back and forth until the solution appeared well-mixed (i.e. not ‘streaky’) and left to sit in the dark at RT for 15 minutes. After transfection, the solution was diluted with 4.8 mL of W5 solution and spun down for 5 minutes at 100 x g for 5 minutes. Transfection solution was removed by pipetting, then pellet was resuspended in 4 mL WI solution (0.5M mannitol, 4mM KCI, 4mM MES (pH 5.7)) and transferred to a 6-well culture plate. After 18 hours of dark incubation at RT, cells were placed under benchtop light (∼25 μmol m^−2^ s^−1^) for 4 hours before RNA harvest.

For all experiments, a GFP-expressing pUC19 plasmid control was used to ensure suitable transfection efficiency and cell health. Any seed batch that was not >80% fluorescent when transfected with GFP was discarded. All library experiments were performed with at least two biological replicates, achieved by running different batches of WT seed on different days. During all steps of the process, seedlings and leaf strips were exposed to as little light as possible to minimize etiolation. As cells are highly sensitive, wide-O pipette tips were used in every instance that cells were pipetted.

### Rice cultivation and mesophyll protoplast isolation

120 to 300 Oryza sativa cv. Kitaake seeds were dehulled and sterilized (75% for 1 minute, 2.5% sodium hypochlorite for 20 minutes, and washed four times with 50 mL ddH_2_O) and planted on 7% ½ MS-Agar (adjusted to pH 5.7 with KOH) in a sterile flow hood. The seeds were incubated for 7 days in a growth chamber (150 μmol m^−2^ s^−1^, 12:12 photoperiod, 27°C/25°C, 80% relative humidity) until the first true leaf was close to being fully developed. The seedlings were left in the chamber for 2 more days with the light intensity halved until the second leaf stage, i.e. to where the leaf was close to full expansion. The second leaves were harvested with their base and tip removed and subsequently sliced into ∼0.5 mm strips perpendicular to the vein on a large petri dish. The leaf strips were immediately transferred to another tin-foil covered petri dish with 0.6 M mannitol and 20 mM MES (pH 5.7) to initiate plasmolysis until the rest of the ∼300 leaves were processed. All leaf strips were subjected to 10 minutes of additional in-dark plasmolysis. The mannitol was then drained and the leaf strips were resuspended in 200 mL of freshly filtered protoplast digestive solution (10 mM KCl, 0.6 M Mannitol, 20 mM MES, 1.5% w/v cellulase (Onozuka R-10), 0.75% w/v macrozyme (Duchefa Biochmi), 0.1% w/v BSA, 10 mM CaCl_2_, 5 mM beta-mercaptoethanol, 50 ug/mL carbinecillin) prior to transfer into 500 mL flasks covered with foil to exclude light. The leaf strips were then subjected to 3 rounds of vacuum infiltration (15 inHg vacuum for 10 minutes each) in 500 mL flasks in the dark and transferred to a dark shaking incubator for digestion of 6 hours (60 rpm for the first 1.5 hours, 70 rpm for the subsequent 3.5 hours and 80 rpm for the last hour prior to harvesting). Mesophyll protoplasts were harvested in two sequential steps. In the initial harvest, a volume of W5 equal to that of the digestive solution was added to quench the digestion reaction. The mixture was then filtered through 40 μm cell strainers into 50 mL conical tubes and centrifuged at 4°C at 200x g for 5 minutes. Green mesophyll pellets were consolidated in W5 as the initial harvest. The second harvest was performed by adding 100 mL of W5 to the leaf strips, swirling and incubating the mixture for 10 minutes. The mixture was then filtered through 40 μm cell strainers into 50 mL conical tubes and spun down at 200x g for 3 minutes. Both harvests were let to sediment naturally under gravity for 1 hour in the dark. The naturally sedimented pellet was isolated and resuspended in 15 mL of MMG. The first and second harvests were pooled followed by viability staining with Evan’s Blue dye. Any preparation that had more than 15% stained cells was discarded. The same transfection protocol used for sorghum protoplasts was used for rice library transfections. Rice protoplasts were exposed to light 12 hours after transfection and harvested for RNA extraction 16 hours post transfection.

### RNA extraction, reverse transcription and library amplification

RNA was extracted using the Qiagen RNeasy Plant Mini Kit (74904) according to manufacturer protocol with the exception of the precipitation step where 1 volume (instead of 0.5 volume) was used for RNA precipitation. On-column DNase digestion was performed according to manufacturer protocol to remove DNA contamination, and 40 uL of DEPC water was used to elute concentrated RNA. Any samples with an A260/230 value below 1.80 were cleaned up using the Monarch® Spin RNA Cleanup Kit (T2030L). Reverse transcription with the Omniscript RT Kit was performed using a 19-mer oligodT primer. Six separate reactions were set up with 1ug of RNA as input. For each sample, cDNA derived from a total of 6ug of RNA was cleaned up using the ZYMO DNA Clean & Concentrator-5 kit with 7:1 DNA binding buffer:cDNA ratio, and used as template for illumina PCR amplification. Gene-specific primers with Illumina PCR1 adaptors were used to amplify barcoded fragments using Q5 HiFi polymerase with the minimal number of cycles to obtain an gel-extractable band (typically 15-20 cycles, primer IDs p219-222). These PCR1 amplicons were gel extracted, cleaned up, amplified to PCR2 products, and sequenced on an Illumina Nextseq 2000.

### Dual-luciferase assay

For the dual-luciferase assay in sorghum, select mutants of the *PsbS* and *SBPase* promoter/5’ UTR region were cloned upstream of a nanoluciferase gene on a pUC19 backbone (Strains). This was performed by producing a WT promoter version of both genes, then introducing mutations via around-the-horn PCR and Gibson homology cloning. The annotated 3’ UTR for each gene tested was attached downstream of the enzyme sequence. Plasmids were constructed to harbor a firefly luciferase gene with a switchgrass Ubi2 promoter and pea RbcS E9 terminator for strong expression.

For sorghum protoplast transfections, at least two biological replicates were used, with a technical replicate for each. One million protoplasts in 1 mL of MMG solution were transfected with 20 μg of plasmid in 40 μL of ddH_2_O in 15 mL conical tubes. Transformation was performed with PEG solution as described above, after which cells were incubated in the dark in 2 mL WI solution for 16 hours and then exposed to diffuse benchtop light (15-20 μmol m^−2^ s^−1^) for 4 hours. For rice protoplast transfections, four technical replicates were performed with similar cell and DNA conditions to the sorghum assay. After transformation, cells were incubated for 12 hours in 2 mL WI solution, then exposed to benchtop light for 4 hours.

For luminescence measurements in both species, cells were spun down in 15mL conical tubes (250x g, 5 minutes, RT), resuspended in 1mL WI buffer, spun down again under the same conditions, then resuspended in 1X passive lysis buffer (Promega). Luciferase and nanoluciferase activity were measured on a Tecan Infinite M1000 Pro using the Promega Nano-Glo Dual-Luciferase Reporter Assay System (N1620) according to the manufacturer’s instructions. Specifically, 80 μL of lysate was combined with 80 μL of luciferase substrate then shaken in the plate reader (orbital, 2 mm diameter, ∼300rpm) then left to sit for 2 minutes before measuring luminescence. The sample was thoroughly mixed with 80μL NanoDLR Stop&Glo Reagent, then shaken (orbital, 1mm diameter, ∼600 rpm) for 3 minutes and left to sit for 12 minutes before measuring nanoluciferase luminescence. Reported values of nanoluciferase/luciferase intensity were normalized to that of an unmodified WT promoter tested in the relevant biological replicate.

### Promoter and Gene Expression Analysis

For tissue-specific expression of candidate genes (Supplementary Fig. 1e), expression values were compiled from the GeneAtlas v2 database^62^ for Sorghum bicolor v3.1.1 for mature leaves (leaf middle whorl.vegetative), seedling leaves (lower leaf upper.juvenile), roots (root top.juvenile), and grain (seed dry grain maturity). For single-cell expression analysis, cell type-resolved RNA-seq profiles from sorghum plants undergoing de-etiolation were taken from a previous study^36^ and queried for target genes. The average expression level of and percent of cells expressing *PsbS*, *Raf1*, and *SBPase* were plotted using the DotPlot function in Seurat version 5.3.0^63^. For single-cell ATACseq analysis, cell type-resolved chromatin accessibility profiles were obtained from a previous study^64^, visualized using the Plant Epigenome Browser^65^, and exported for combined visualization in Geneious Prime 2025.2.2. The PlantCARE ^66^ and JASPAR databases ^67^ were used for annotating putative TFBSs and identifying high-effect cREs.

For local sequence alignments in the Poaceae, mVISTA was used^68^. Promoter (defined as the 2 kb upstream from the annotated transcription start site) and 5’ UTR sequences for each gene were retrieved from the Phytozome database for sorghum (Sorghum bicolor RTx430 v2.1), maize (Zea mays RefGen_V4), miscanthus (Miscanthus sinensis v7.1), *Setaria italica* (Setaria italica v2.2), brachypodium (Brachypodium distachyon v3.2), and rice (Oryza sativa v7.0). Sequences were aligned using mVISTA with alignment windows of 100 bp at a similarity threshold of 70% and plotted between 50% and 100% sequence similarity. For base-pair resolved sequence alignments, Clustal Omega was used via the EMBL-EBI portal^69^.

For displaying single-nucleotide variant effects, we generated a position-specific sequence logo that visualizes substitution preference at each genomic position based on measured effect sizes. For each position, we converted the base-specific effect sizes into a probability distribution (*P_i_* at position i) via a softmax transform with temperature parameter β_scale (set to 5). We then computed the information content per position as *IC_i_* = 2 - *H(P_i_)*, where the term *H(P_i_)* quantifies the Shannon entropy (bits; maximum IC = 2 for DNA). The plotted letter heights are *P_b,i_ x IC_i_,* where *P_b,i_* is the softmax-normalized preference for base *b* at position *i*. Logos were rendered with logomaker using a colorblind-aware palette.

### Library Design

Using gene models from Phytozome *Sorghum bicolor* RTx430 v2.1 (PsbS gene ID: SbiRTX430.03G398900, Raf1 gene ID: SbiRTX430.06G286900, SBPase gene ID: SbiRTX430.03G387300), we designed libraries to span −2000 bp upstream of the annotated transcriptional start site (TSS) until the beginning of the protein coding sequence, i.e. including the 5’ untranslated region. The designed libraries include 4x 500 base pair deletions, 10x 200 base pair deletions for the entire region. In the region 1 kilobase of the transcription start site through the UTR all single nucleotide substitutions, all 2 base pair deletions, all 12 base pair deletions and A-to-G transitions within each 5 base pair window were designed. In addition, we inserted putative transcription factor motifs every 5 base pairs in the forward and reverse direction at positions −150 to +50 from the transcriptional start site. We chose to insert 40 motifs from each source: 1, imputed transcription factor motifs catalogued in PlantTFDB^18^ for Sorghum leaf tissue expressed transcription factors identified in the Sorghum Riken Database (http://sorghum.riken.jp/), and 2, DNase I footprints identified in sorghum^19^. Details of library design can be found on github (https://github.com/SavageLab/plant_promoter_bashing).

### Library Cloning

We performed all cloning using a pUC19 plasmid backbone containing an ampicillin resistance cassette and TOP10 *E. coli* cloning strain. Cells were grown in Luria-Bertani broth supplemented with 0.1mg/ml carbenicillin at 37°C. All polymerase chain reactions (PCRs) were performed using a Bio-Rad T100 Thermal Cycler. All PCR products were purified using Zymo DNA Clean & Concentrator-Kit (D4033) after amplification, and before electroporations. All library cloning steps were performed *in vitro* until the final bottleneck step in order to prevent bias to the library.

We first created wild type plasmids for each gene (Extended Data Fig. 3a). In the case of the synthetic construct, this contained the gene promoter/5’ UTR driving GFP with a pea rcS E9 terminator. In the case of the natural gene construct, plasmids contained the full gene including 2 kilobases of promoter, 5’ UTR, all introns and exons, and an additional 1 kb downstream to capture the native 3’ UTR sequence. To do so, we used primers (primer IDs p1-p16) to amplify the promoter or full length gene from Sorghum RTx430 genomic DNA using NEBiolabs Q5^®^ High-Fidelity 2X Master Mix (3 minutes at 95°C; 35 cycles: 20 seconds at 98°C, 20 seconds at 67°C, 2-3 minutes at 72°C; 1 minute at 72°C). We performed around-the-horn PCR (Primer IDs p223-p224) using KAPA HiFi HotStart ReadyMix (Roche: 09420398001) (3 minutes at 95°C; 35 cycles: 20 seconds at 95°C, 20 seconds at 66.4°C, 15 seconds at 72°C; 1 minute at 72°C) to linearize the pUC19 destination plasmid. These fragments were then combined using Gibson assembly (NEBuilder HiFi DNA Assembly Master Mix (E2621L) (60 minutes at 50°C). These Gibson reactions were transformed into chemically competent TOP10 cells, and individual clones were isolated and sequence verified.

Next, we cloned a random barcode into each plasmid. To do so, we performed an around the horn PCR using primers containing randomized 15 base pairs (Primer IDs p213-p218 for native libraries, p225-p226 for GFP libraries), and then used a selfing Gibson reaction to circularize our plasmids. These randomized plasmid pools then served as the template to introduce the promoter and 5’ UTR variants. An unused subsample of these reactions were transformed into electrocompetent TOP10 cells to ensure a sufficient amount of barcodes were obtained via titer plating, and Illumina sequencing was performed (Illumina iSeq, Kit v2) on the barcoded region to ensure barcode diversity and count was suitable for downstream cloning.

For constructing the large deletions (200 and 500 base pair deletions), we performed an around the horn PCR using primers that miss these specific regions (primer IDs p17-p114) with KAPA HiFi HotStart ReadyMix (Roche: 09420398001; 3 minutes at 95°C; 35x: 20 seconds at 98°C, 20 seconds at 65°C, 5 minutes at 72°C; 5 minutes at 72C and hold at 4°C) on the respective gene-specific wild type plasmid. Linearized DNA was then circularized using Golden Gate reactions with BsaI-HF-v2 (NEB: R3733S) for *Raf1*, BsmBI-v2 (NEB: R0739S) for *SBPase*, and AarI (NEB: R0745S) for *PsbS* (for BsaI and AarI using 30 cycles of 5 minutes each at 37°C and 16°C; for BsmBI using 30 cycles of 5 minutes each at 42°C and 16°C).

To create the higher resolution variant libraries, we ordered ∼270 base pair oligopools from Twist Bioscience. Since the mutagenized region (>1 kilobase) is larger than the synthesis length limit, we split the oligopool into 7 to 8 different 200 base pair fragments, and performed individual Golden Gate assemblies for each fragment and gene combination. Each gene sublibrary was amplified using KAPA HiFi HotStart ReadyMix (Roche: 09420398001) from the PCR cleaned synthesized pool (using primers p159-p202: 3 minutes at 95°C; 35 cycles: 20 seconds at 98°C, 15 seconds at 57°C, 15 seconds at 72°C; 1 minute at 72°C). Corresponding Golden Gate compatible linearized vectors were amplified using KAPA HiFi polymerase (using primers: p115–158: 3 minutes at 95°C; 35 cycles: 20 seconds at 98°C, 15 seconds at 65°C, 1 min/kb at 72°C; 10 minutes at 72°C).

We created synthetic construct libraries first with all desired mutations in the plasmid pool, and then amplified and ported the entire promoter/5’ UTR region into the barcoded natural gene constructs. Full promoter/UTR library regions were amplified from the synthetic libraries alongside the conjugate barcoded backbone vectors (using Q5 2X Mastermix; primers p203-212; using 3 minutes at 95°C, 35 cycles of: 20 seconds at 98°C, 20 seconds at 65°C, 5 minutes at 72°C; 5 minutes at 72C and hold at 4°C) and circularized using Gibson Assembly (NEBuilder HiFi DNA Assembly Master Mix (E2621L), 60 minutes at 50°C).

We then PCR purified the assembly products, and electroporated 50 ng into 50ul of electrocompetent TOP10 cells using 0.1cm cuvettes (Bio Rad 1652089) using a Bio Rad Micropulser using the EC1 setting (1.8 kV). Cells were grown at 37°C to OD∼0.5 to limit the number of cell divisions causing library bias, and cryostocks were made by 1:1 dilution with 50% glycerol. Cells were then bottlenecked to around 100,000 clones by plating on LB/carbenicillin agar plates to limit the maximum number of individual barcodes propagated. All library construction steps until the final bottlenecking were performed *in vitro* to minimize mutant or barcode bias.

### Long read sequencing for mapping barcodes to promoter variants

In order to link individual barcodes with the promoter variants, we performed long read sequencing using PacBio Revio. To do so, we first grouped reads by their barcode, and created a consensus sequence for each barcode using samtools v1.2 (using ‘consensus –config hifi’). We then mapped consensus sequences to the wild type reference plasmid using minimap2 v2.26-r1175 (using ‘–MD-Lax map hifi’) to identify mutations in each consensus read. All consensus variants were length filtered to be no less than 600 bases shorter than the wild type plasmid. We observed 100,883 SBPase, 33,126 Raf1 and 47,423 PsbS barcode-to-variant mappings.

### Short read, high volume sequencing to determine expression levels for barcodes

To determine the expression changes due to a crDNA mutation, we used deep Illumina sequencing of the plasmid library, as well as cDNA isolated from RNA after 16-18 hours post protoplast transformation. For illumina sequencing analysis we followed the following workflow: We merged paired end reads using FLASH (v1.2.11), with a maximum number of overlap bases of 150. We then applied a quality filter using vsearch (v.2.28.1) (using - fastq_truncqual 20, –fastq_maxns 3, fastq_max33 0.5). This allowed us to count and calculate a log_2_ read ratio post-expression vs. pre-transformation for each barcode using a pseudocount of 1. We then merged these barcode read ratios with promoter variants identified by long read sequencing. We then removed any barcodes that have mutations in priming site for barcode sequencing (for SBPase: positions 1280-1335 and 1448-1484, for Raf1: 675-710 and 485-525, for PsbS 1360-1393 and 1192-1211 relative to the reference plasmid sequence), since we detected that mutations here introduce artefactual skew on the log read ratios independent of the crDNA variants observed. For Raf1 we also observed mutations “T3575A”, “C5576T”, “A5574G” and “T5575A” (relative to the reference plasmid sequence) in almost all reads, and redefined these mutations as the new reference sequence.

### Individual Mutant Effect Inference

The majority of reads contained multiple mutations due to synthesis, cloning and sequencing artefacts. These are particularly pronounced at poly-nucleotide repeats, where addition or subtraction of one repeat-base is observed frequently (Extended Data Fig. 2a). For example, for PsbS, we find that >80% of reads contain an additional A base pair insertion at a 10-base pair poly-A tract 297 bases downstream from the translational stop site. To deconvolute potentially casual individual variant effects, we performed ordinary least squares linear regression using statsmodels v0.14.5. Using additive assumptions, this enables inference of whether a particular individual mutation can explain the observed expression effects additionally to co-occurring mutations, including sequencing and cloning artefacts, to pinpoint potential causal mutations (Extended Data Fig. 2c). For example, a mutation that co-occur in almost all reads, such as the poly-nucleotide repeat mutations, cannot explain overexpression in a subset of high-expressing combinations of mutations. We stringently called variants significant and meaningful if their p-value passes a 0.05 Bonferroni correction, and if both their inferred effect as well as the mean log read ratio of reads containing that mutation is larger than 1.5. We included this raw log read ratio cutoff in order to prevent erroneously negatively called passenger mutants from driving a co-occurring candidate mutant’s positive inferred effect without having observed that candidate mutant in hypermorphic reads.

### Fine tuning the genomic language model

We developed an automated pipeline to create a gene expression prediction dataset from RNA-seq data in Expression Atlas^70^. The pipeline is available on github (https://github.com/songlab-cal/gpn/tree/main/workflow/make_expression_dataset_from_gxa) and should be applicable to many other species available in Expression Atlas. As input, the user must specify a list of experiment accessions. In our case we chose Sorghum experiments using the reference cultivar BTx623: E-MTAB-4021, E-GEOD-98817, E-MTAB-4203, E-MTAB-4273, E-MTAB-4400, E-CURD-25, and E-MTAB-5956. The pipeline has the following steps: 1. Download transcript-level transcript per million (TPM) values. 2. Filter samples with variance less than a threshold (default: 1). 3. Drop conditions with no biological replicates, or those where Pearson correlation between replicates is less than a threshold (default: 0.8). 4. Average expression among replicates. 5. Apply a log1p transformation, so the final prediction target is log(1+TPM). 6. For each transcript, extract the reference genome sequence around the TSS (default: +-256bp).

This resulted in a total of 26 conditions, found under https://huggingface.co/datasets/gonzalobenegas/gxa-sorghum-v1/blob/main/labels.txt. GPN-Brassicales^45^ was finetuned in a multi-task fashion to predict all of the conditions, using a mean-squared error (MSE) loss. To evaluate on MPRA data, the condition “E-MTAB-4021_leaf mesophyll” was used. The model architecture was the following: 1. GPN base pair-level embeddings from the last layer were first averaged across spatial positions. 2. An MLP mapped this high-dimensional embedding into pre-activations for each output condition. 3. Pre-activations were transformed into positive predicted gene expression values using the softplus activations.

The model was trained on chromosomes 1-8. Chromosomes 9-10 were left for potential validation and testing, though we did not find held-out performance on RNA-seq useful for indicating performance on MPRA variants. The following hyperparameters were used, chosen to be reasonable defaults but not systematically tuned: Max epochs: 30, Batch size: 128, Optimizer: AdamW, Learning rate: max 1e-3, with 1% warmup ratio, cosine decay down to 1e-4, Weight decay: 0.01.

## Supporting information

primers and strains

## Acknowledgments

We thank Christine Queitsch and Josh Cuperus for experimental guidance and comments on the manuscript. We thank Jen Sheen, Tobias Jores, Cole Mueth, and Sayeh Gorjifard for their advice and guidance in developing a sorghum MPRA assay, and Stan Fields, Nicholas Karavolias, Dhruv Patel-Tupper, Tamara Miller, and Luke Oltrogge for their essential help in formal analysis and in building the scientific narrative. We thank Jianqiang Shen, Sultana Anwar, and Kiflom Aregawi for providing sorghum seed, Armen Kelikian and Amala John for their consultation on nonphotochemical quenching biology and *PsbS* natural diversity, and Netra Krishnappa and Shana McDevitt for assistance in high-throughput sequencing. This article is subject to HHMI’s Open Access to Publications policy. HHMI lab heads have previously granted a nonexclusive CC BY 4.0 license to the public and a sublicensable license to HHMI in their research articles. Pursuant to those licenses, the author-accepted manuscript of this article can be made freely available under a CC BY 4.0 license immediately upon publication.

## Author contributions

Conceptualization: EDG, DD, FZW, BJS, KKN, DFS.

Methodology: EDG, DD, FZW, GB, KKN, YSS, DFS.

Software: DD, GB, EDG, SC, MM, YSS.

Validation: EDG, FZW, VG.

Formal analysis: DD, EDG, GB, YSS.

Investigation: EDG, DD, FZW, GB, JR, SS, MM, VG.

Writing - Original Draft: EDG, DD, DFS.

Writing - Review & Editing: all authors Visualization: EDG, DD, GB.

Supervision: BJS, KKN, YSS, DFS.

Project administration and funding acquisition: DFS.

## Data and materials availability

Generated plasmids are available upon request. Raw sequencing reads can be found at the NCBI Sequencing Read Archive (accession: PRJNA1308589). Analysis pipelines, visualization notebooks, and inferred mutation effects can be found on github (https://github.com/SavageLab/plant_promoter_bashing). Raw per barcode mutant calls, read counts, and log read ratios can be found under: https://drive.google.com/drive/folders/1il-SFZb29KMPypJzViL72edjR7pvddpN?usp=sharing. Expression dataset curation for GPN training and finetuning code is available on github (https://github.com/songlab-cal/gpn/tree/main/workflow/make_expression_dataset_from_gxa, https://github.com/songlab-cal/gpn/tree/main/analysis/gpn_sorghum_expression). The resulting GPN dataset and model are available at huggingface (https://huggingface.co/collections/songlab/sorghum-gene-expression-prediction-68963dd31658bfb98c07ae1b).

---

**Extended Data Fig. 1.**
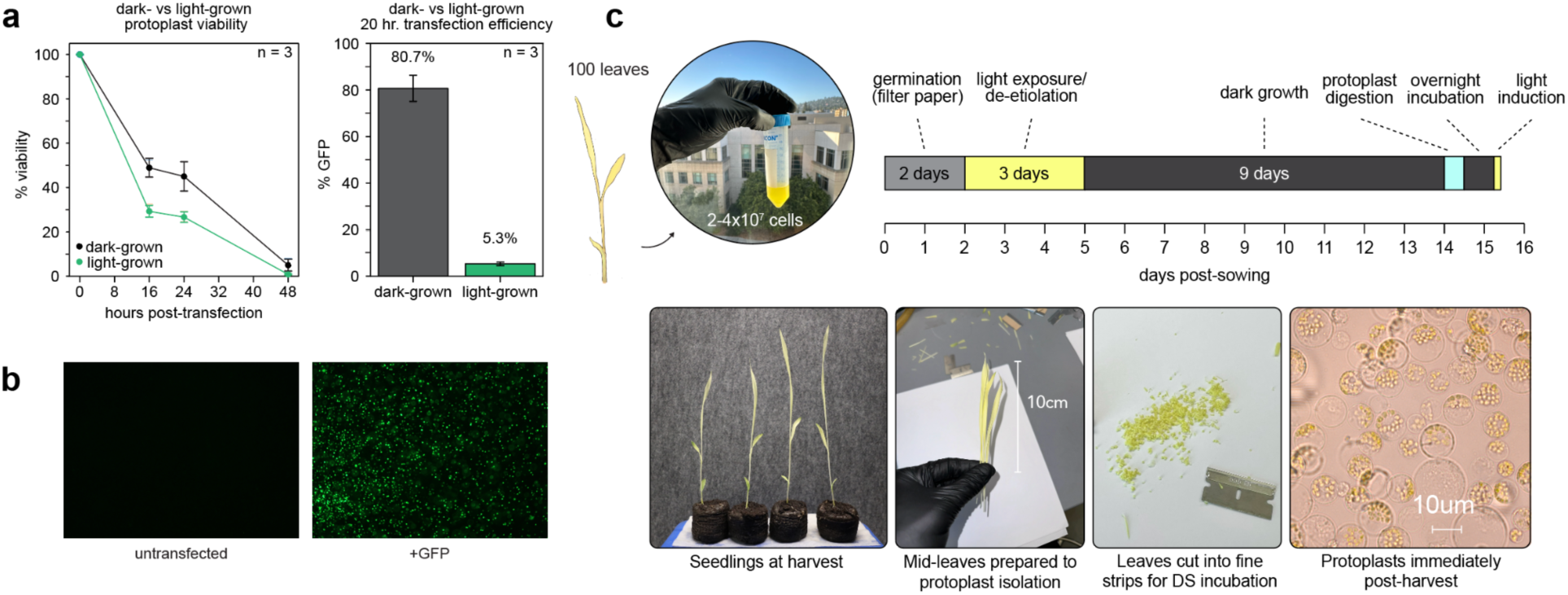
An optimized protocol for high-quality, high-yield protoplasts. **a,** Post-PEG transfection viability of protoplasts isolated from seedlings grown with and without a 9-day dark period before isolation (left). Percent of live cells expressing a GFP reporter 20 hours after harvest is indicated (right). For all measurements, 3 biological replicates (indicating separate seed batches) were used. **b,** Cells after 20 hr of incubation with and without a GFP reporter plasmid transfected. **c,** Image of healthy protoplasts after harvest, indicating the approximate expected yield for 100 seedlings. At right the growth cycle of etiolated seedlings is indicated, alongside timing for optimal harvest. Below, the major steps of protoplast harvest are depicted to emphasize tissue health and leaf-cutting technique.

**Extended Data Fig. 2.**
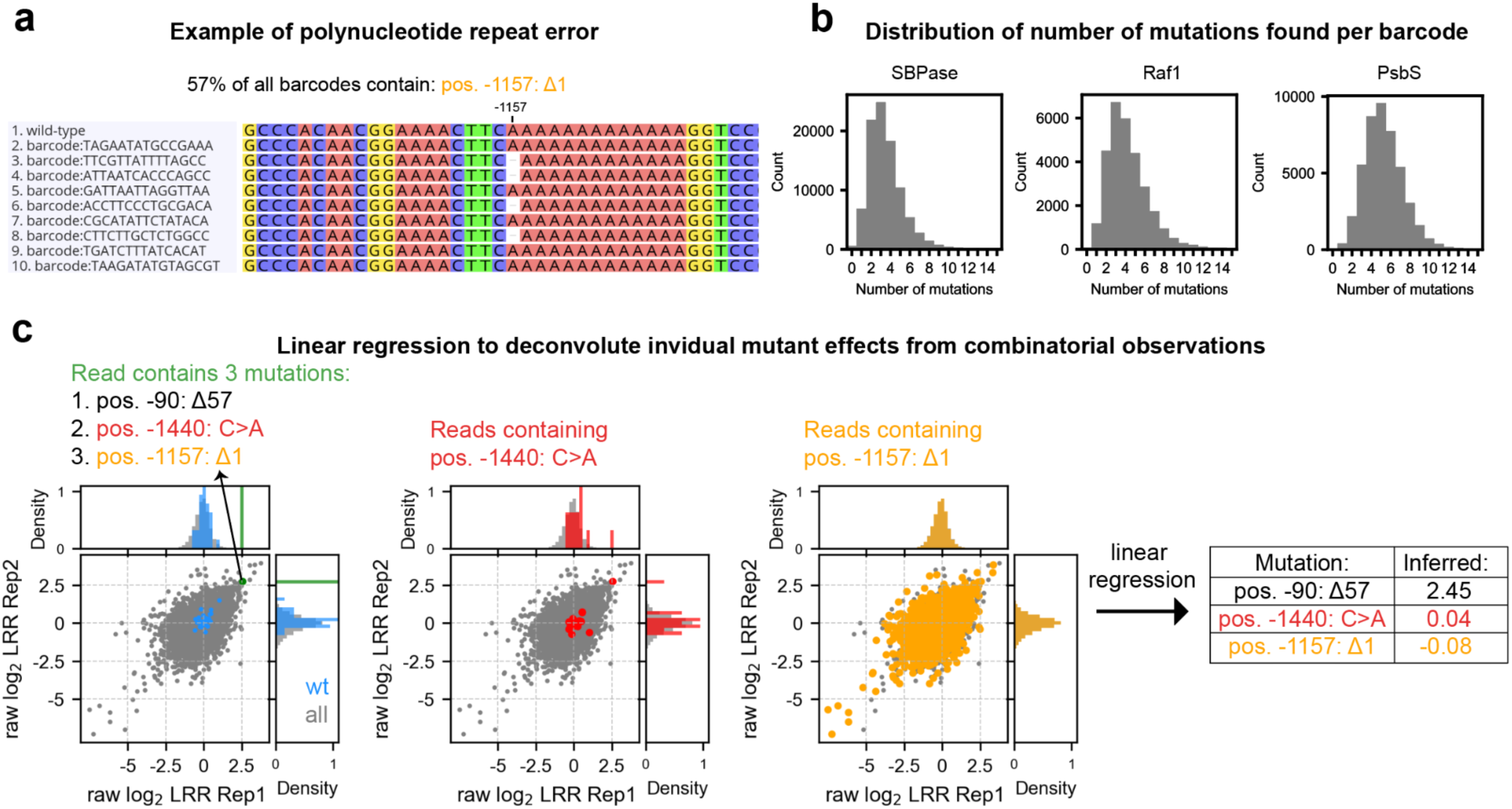
Linear regression enables inference of putative causal individual mutant effects E_mut_ driving changes in expression. **a,** An example of a poly-nucleotide repeat sequencing error. Consensus reads for each barcode are shown aligned to the reference promoter around position −1157 from the TSS of the PsbS promoter. **b,** Errors arising from repeat sequencing, cloning, and synthesis inflate the number of mutations observed per barcode across all samples compared to the designed library (number of mutations per barcode being one) as shown in histograms of the number of mutations per barcode across crDNA libraries. **c,** Scatterplots of log read ratios for each barcode are shown for each biological replicate. A particular high log read ratio barcode (green) contains three mutations (57 bp deletion at pos. −90; C-to-A mutation at pos. −1440; 1bp deletion at pos. −1157). To infer plausible individual mutation effects of each of these three mutations, linear regression considers the additive effect of each individual mutation based on the total dataset to explain the observed combinatorial mutation effect for each barcode. In this example, reads that contain the C-to-A mutation at pos. −1440 (red) and 1bp deletion at pos. −1157 homopolymer error (orange) follow the entire mutation distribution apart from the read in which they are found together with the 57 bp deletion at pos. −90 (left plot, green). This suggests that the 57 bp deletion at pos. −90 drives hypermorphic expression. Linear regression infers a hypermorphic individual effect for the 57 bp deletion at pos. −90, but not for the other two mutations (right table).

**Extended Data Fig. 3.**
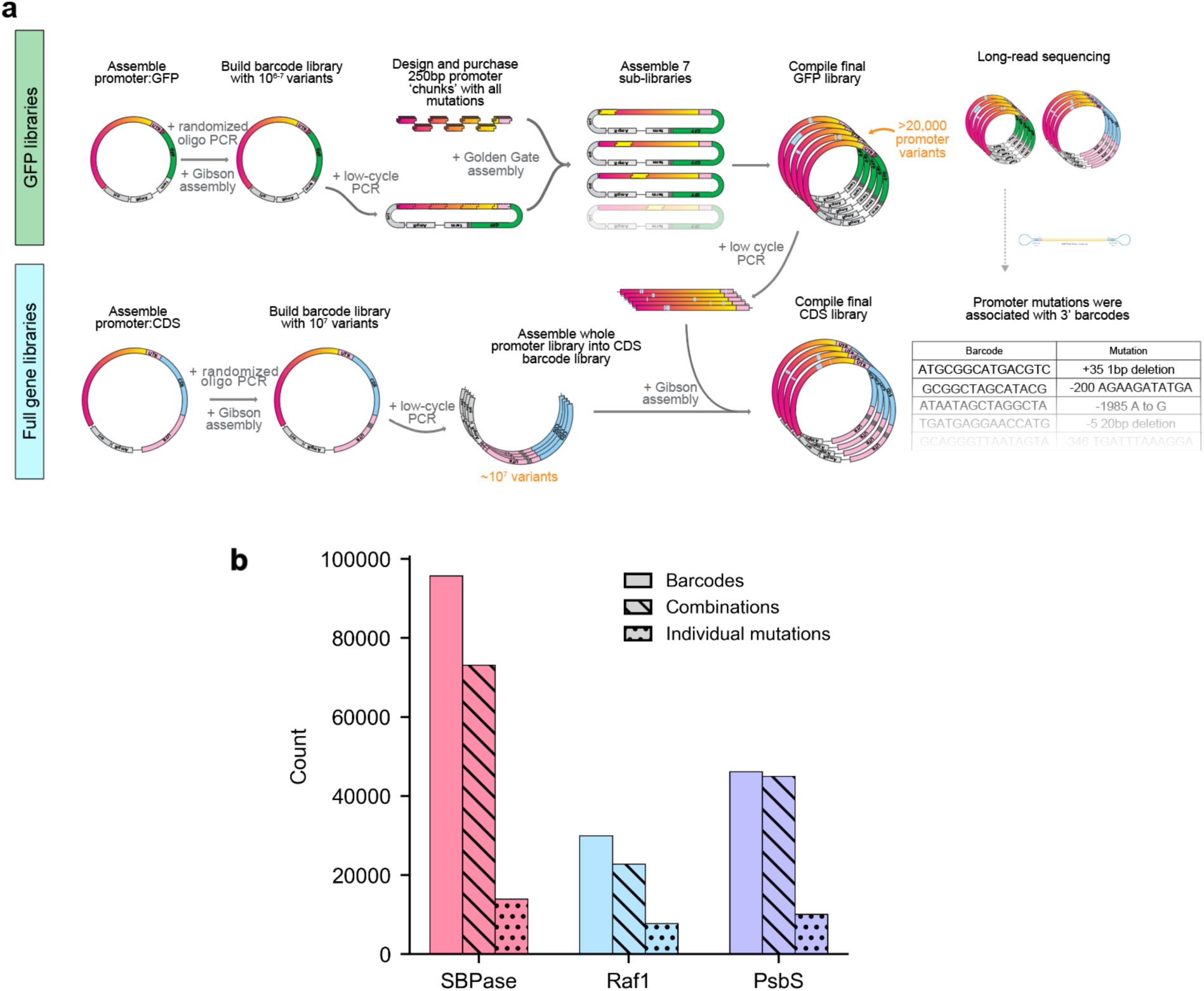
Workflow for generating barcoded variant libraries, and the resulting variant counts are displayed. **a**, Diagram of major steps of library cloning. Libraries were constructed for each gene promoter driving both GFP and the native gene sequence from sorghum. For GFP libraries, promoter and 5’ UTR were cloned into a GFP DNA barcode library with the first 30 bases (10 amino acids) of the native coding sequence fused to GFP. **b**, Barplot displaying the number of unique barcodes, combinatorial variants found among these barcodes, and individual mutations found among the combinations in native gene libraries as determined by PacBio long-read sequencing.

**Extended Data Fig. 4.**
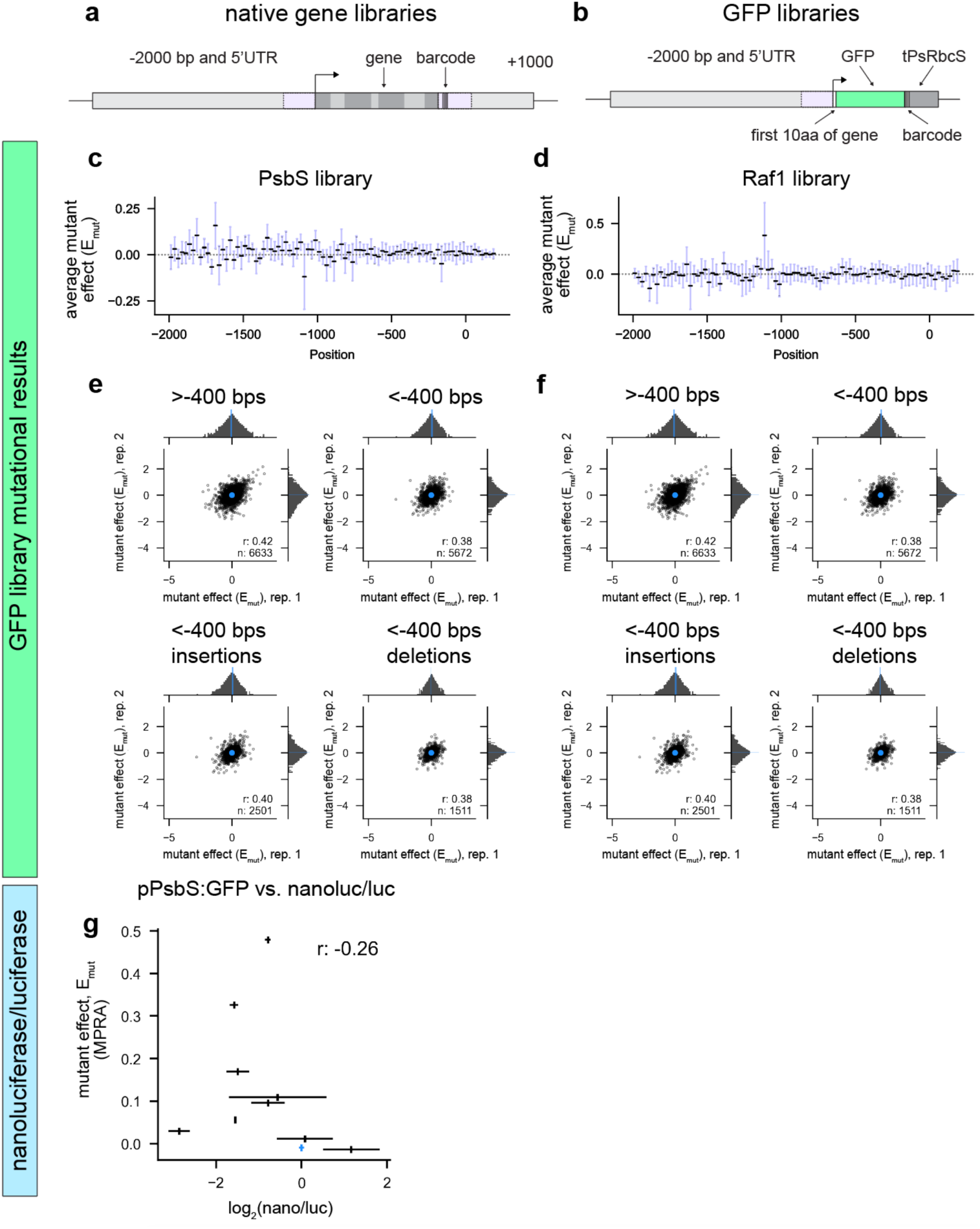
Variant effects measured in a synthetic construct driving GFP-expression neither show reproducible effects nor validate in protein production. **a-b,** Plasmid architectures for native gene plasmids (a) and GFP plasmids (b). For GFP plasmids, GFP is expressed with an N-terminal addition of the first 10 amino acids of the cognate gene and 15 bp DNA barcode is included immediately after the stop codon. GFP is flanked by a 3’ terminator from the *Pisum sativum rbcS* gene. **c-d,** Variant effects measured in GFP-driving constructs do not show any particular hotspots in mutational sensitivity for *PsbS* (c) or *Raf1* (d). **e-f,** Reproducibility between biological replicates for various regions (distal: <-400 bp from transcriptional start site, core: > −400 bp) and mutation types are shown for *PsbS* (**e**) or Raf1 (**f)** promoters driving *GFP* expression. **g,** Comparison of *PsbS* synthetic (GFP) library MPRA enrichment and nanoluciferase/luciferase values for ten promoter/5’ UTR variants. The Pearson’s r is indicated.

**Extended Data Fig. 5.**
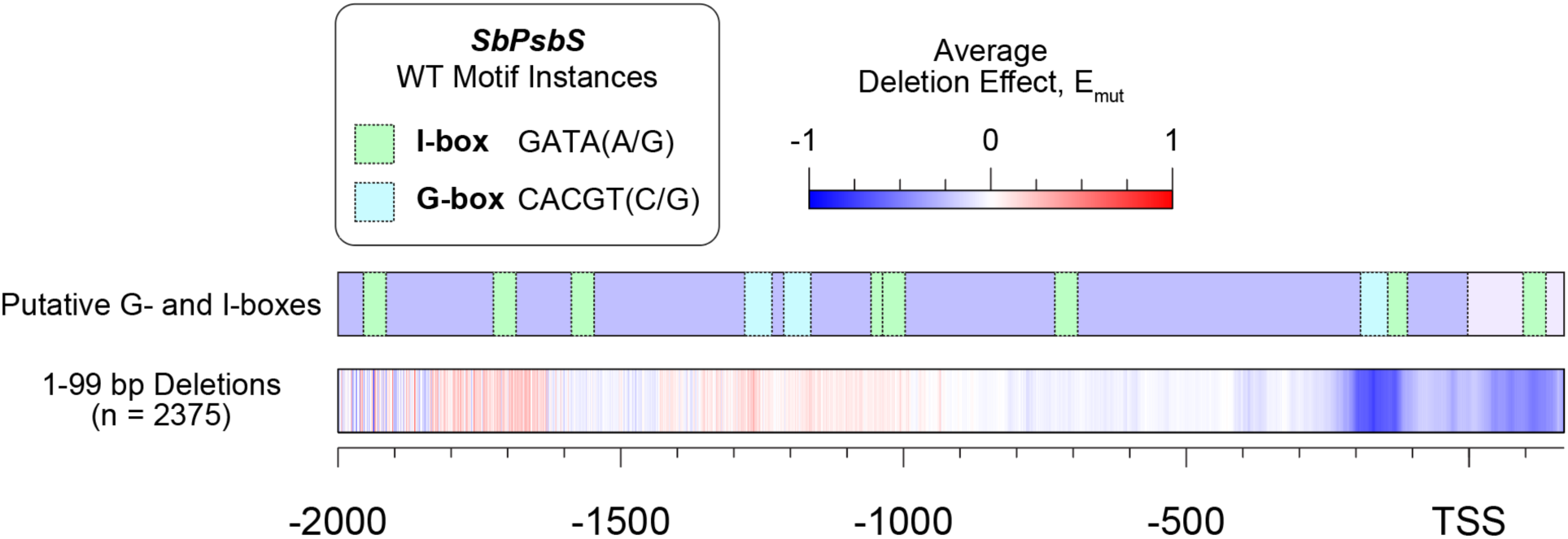
The location of G- and I-boxes in the *PsbS* promoter. Display of all putative G-(cyan) and I-box-like (green) sequences in the *PsbS* promoter and 5’ UTR region based on sequence and predicted TFBS, compared to the average effect of 1-99 bp deletions across the same range.

**Extended Data Fig. 6.**
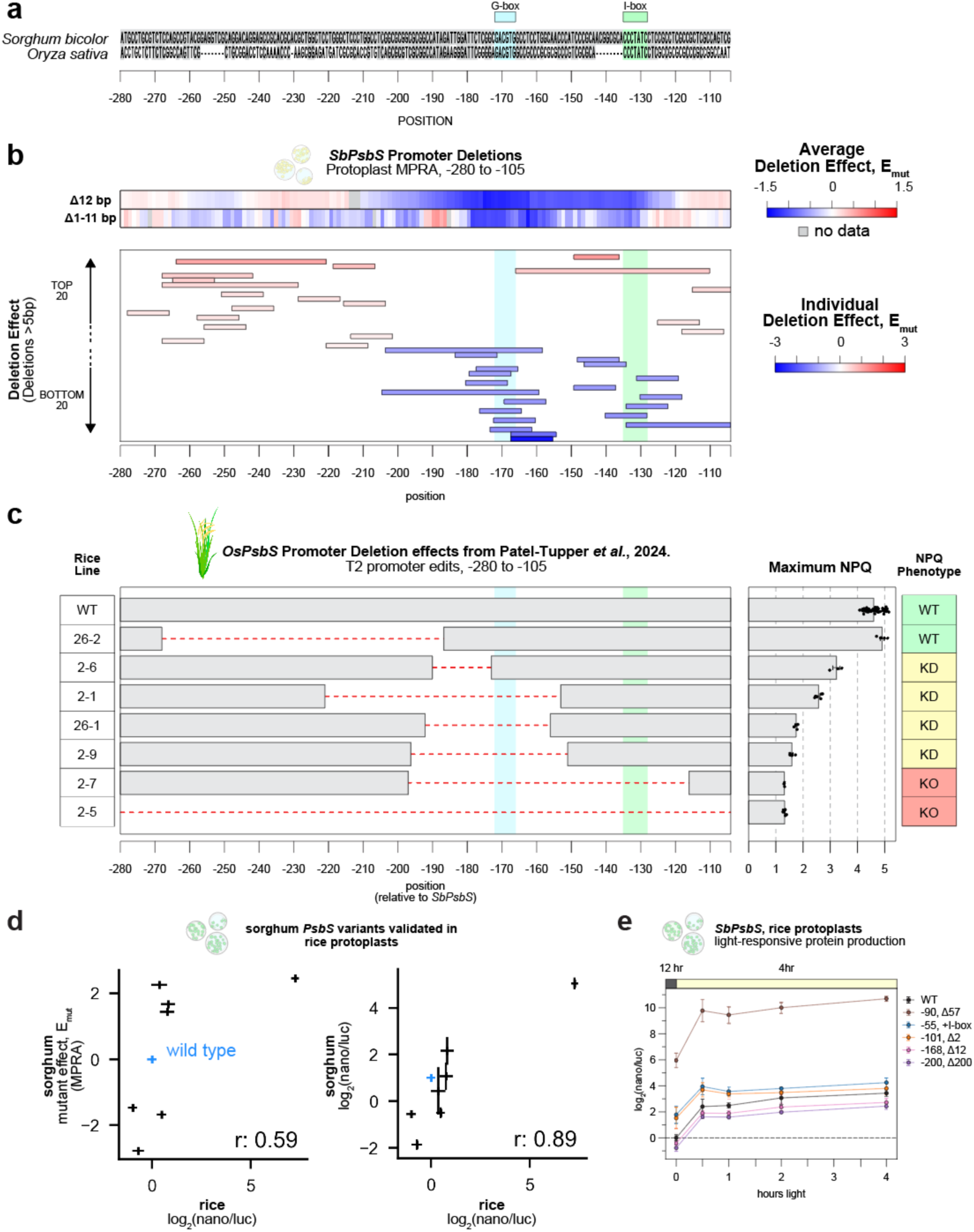
Hypomorphic variants from sorghum MPRA measurements correspond to *in planta* rice quantitative trait engineering results. **a,** Sequence alignment (Clustal Omega) of the sorghum and rice (cv. Nipponbare, ssp. Japonica) *PsbS* promoter in the region −280 to −105 relative to the sorghum TSS. Putative G-(cyan) and I-box (green) locations are indicated. **b**, Measured sorghum MPRA deletion effects averaged for 12 bp and 1-11 bp deletions (top two heatmaps), and individual deletion effects (bottom blot) in the region −280 to −105 are shown. For individual deletions, top 20 hypermorphic and bottom 20 hypomorphic deletions are displayed and colored corresponding to their effect. G- and I-box regions are colored. **c,** Maximum NPQ measurements from T2 non-transgenic *OsPsbS* promoter mutants produced in a previous QTE study^14^. Lines are labelled with their indicated labels and described NPQ phenotype (KD = knockdown, KO = knockout) and deletion position is labelled relative to *SbPsbS* as determined by sequence alignment.

**Extended Data Fig. 7.**
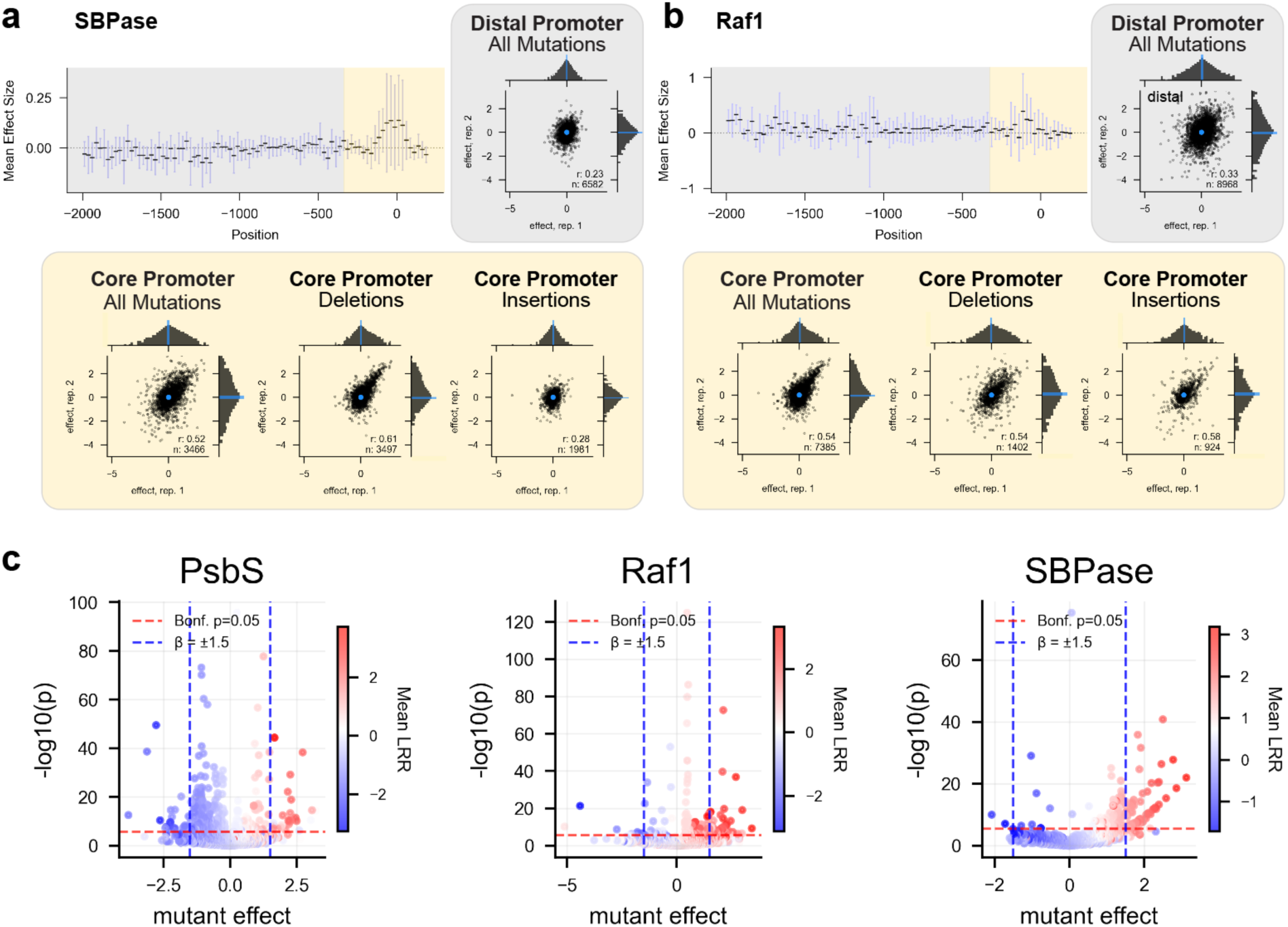
Promoter variant effect measurement across three native gene crDNA libraries reveal significant and meaningful expression modulation. **a-b,** The variance and mean of variant effects across the promoter and 5’ UTR is shown (top left) for *SBPase* (**a**) and *Raf1* (**b**) as measured in their native gene constructs. The correlation between biological replicates within various regions and among different classes of mutations of the promoter are shown for each gene, with the Pearson correlation coefficient, r, and number of observations, n, indicated. **c,** Volcano plots for inferred individual mutant effects are shown, with mutant effects colored by the average log_2_ read ratio of reads they are found in.

**Extended Data Fig. 8.**
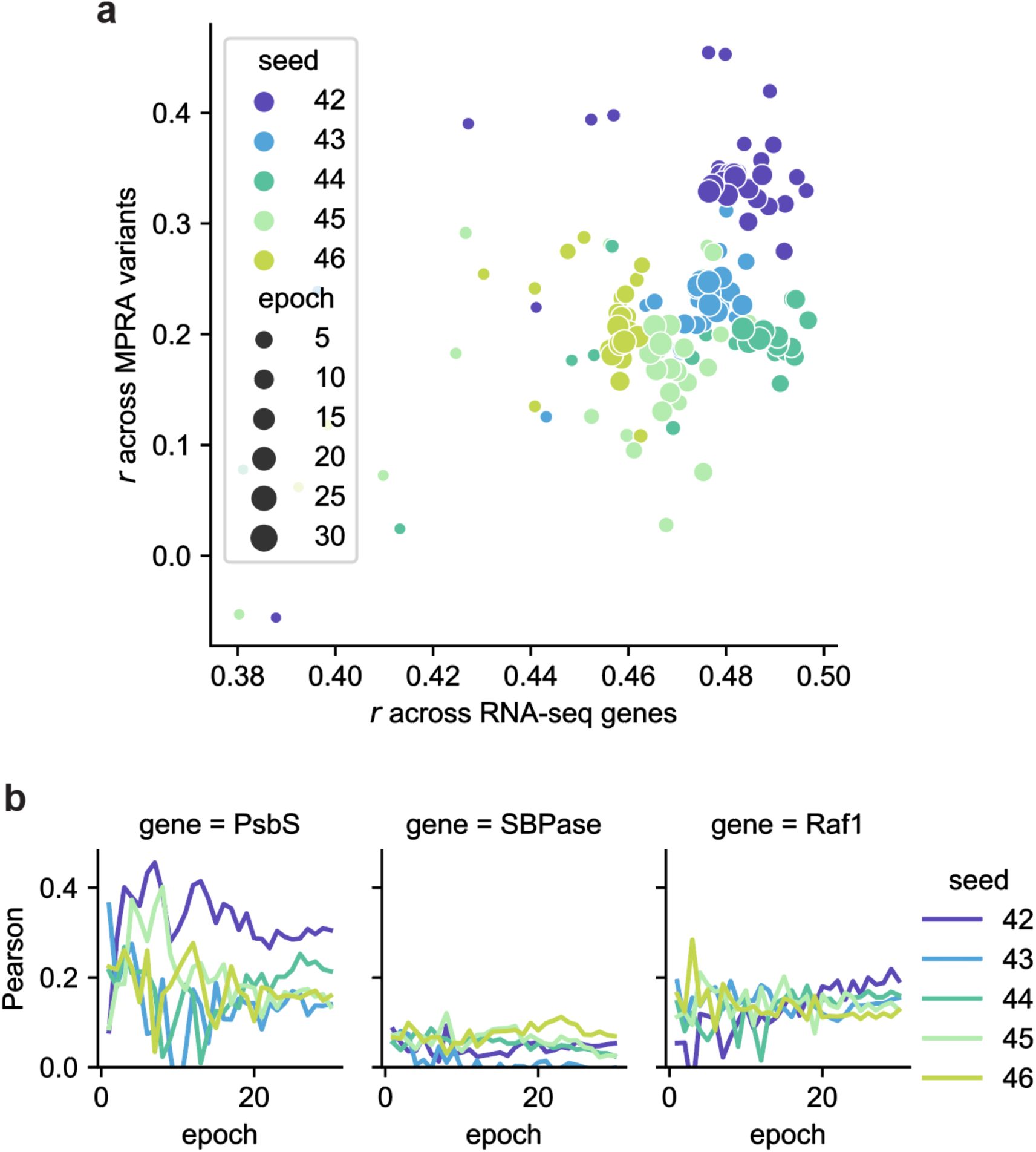
Seed and epoch dependence of GPN-finetuning. **a,** The relationship of model predictions of RNA expression vs. the model predictions across all massively parallel reporter assay (MPRA) measurements in *PsbS* is shown. **b,** The validation set (non-SNP variants) model performance in terms of Pearson r is shown across epoch for each gene and seed.

## Supplementary Figures

**Supplementary Fig. 1.**
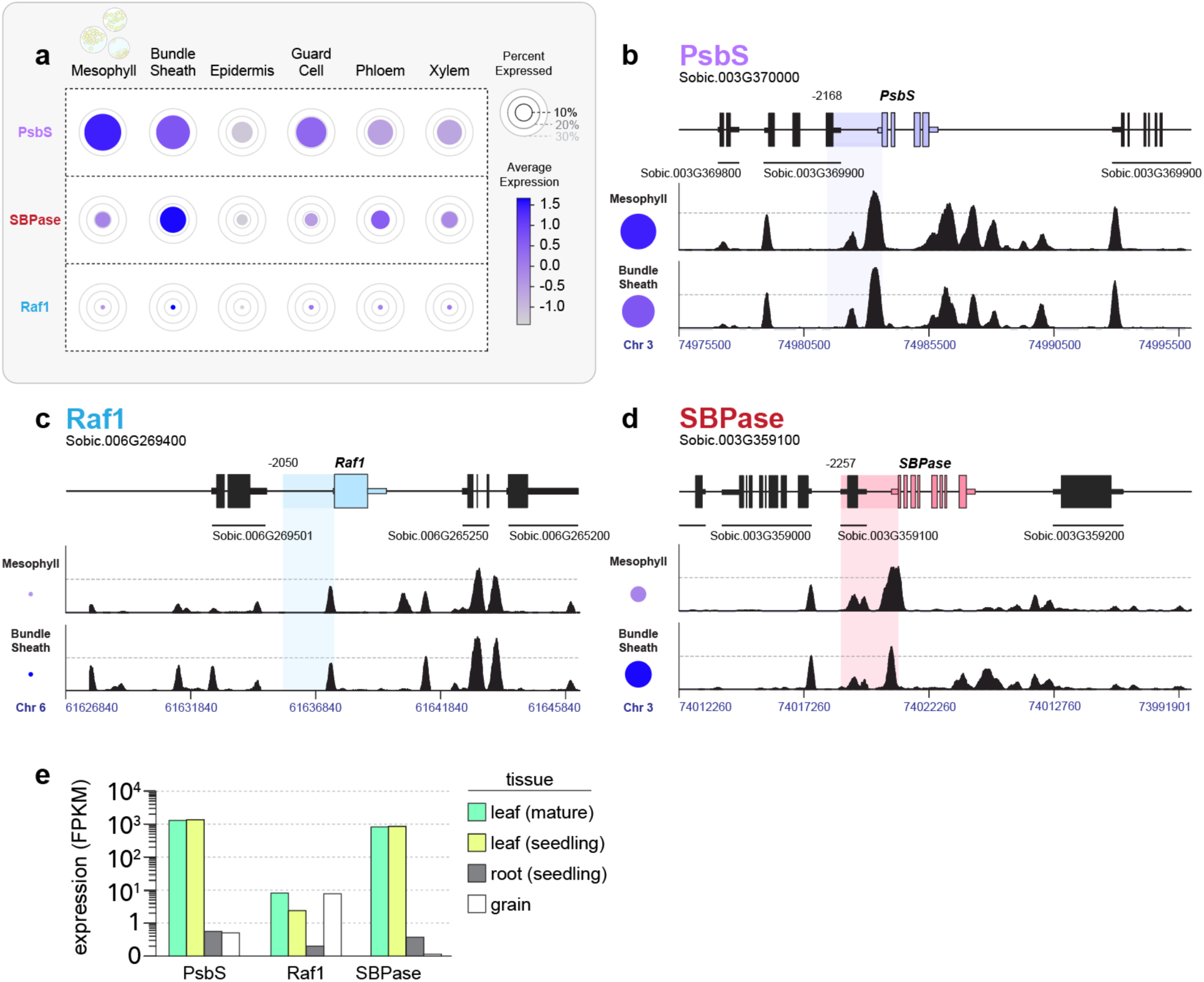
Single-cell profiling of gene expression and chromatin accessibility in rate-limiting photosynthesis genes. **a**, Single-cell RNAseq analysis describing the percentage of cells within a foliar cell niche expressing each gene (circle circumference) and the average log normalized counts scaled to have mean 0 and variance 1 within each cell population (colormap)^36^. **b-d,** Single-cell ATACseq peaks for the ∼20kb window containing each gene isolated from third leaf BTx623 sorghum seedlings^64^. Gene name and locus tag is indicated followed by the location and locus tag of all surrounding genes. Each colored region corresponds with the promoter and 5’ UTR regions sampled for promoter libraries with its distal bound relative to the start codon labelled. For each gene, mesophyll and bundle sheath ACRs are indicated with a common axis (au) with genomic position indicated. **e**, Tissue-level expression profiles (FPKM) for the three surveyed genes collected from the JGI GeneAtlas v2 database^62^. For information on tissue type and sampling see Methods, Promoter and Gene Expression Analysis.

**Supplementary Fig. 2.**
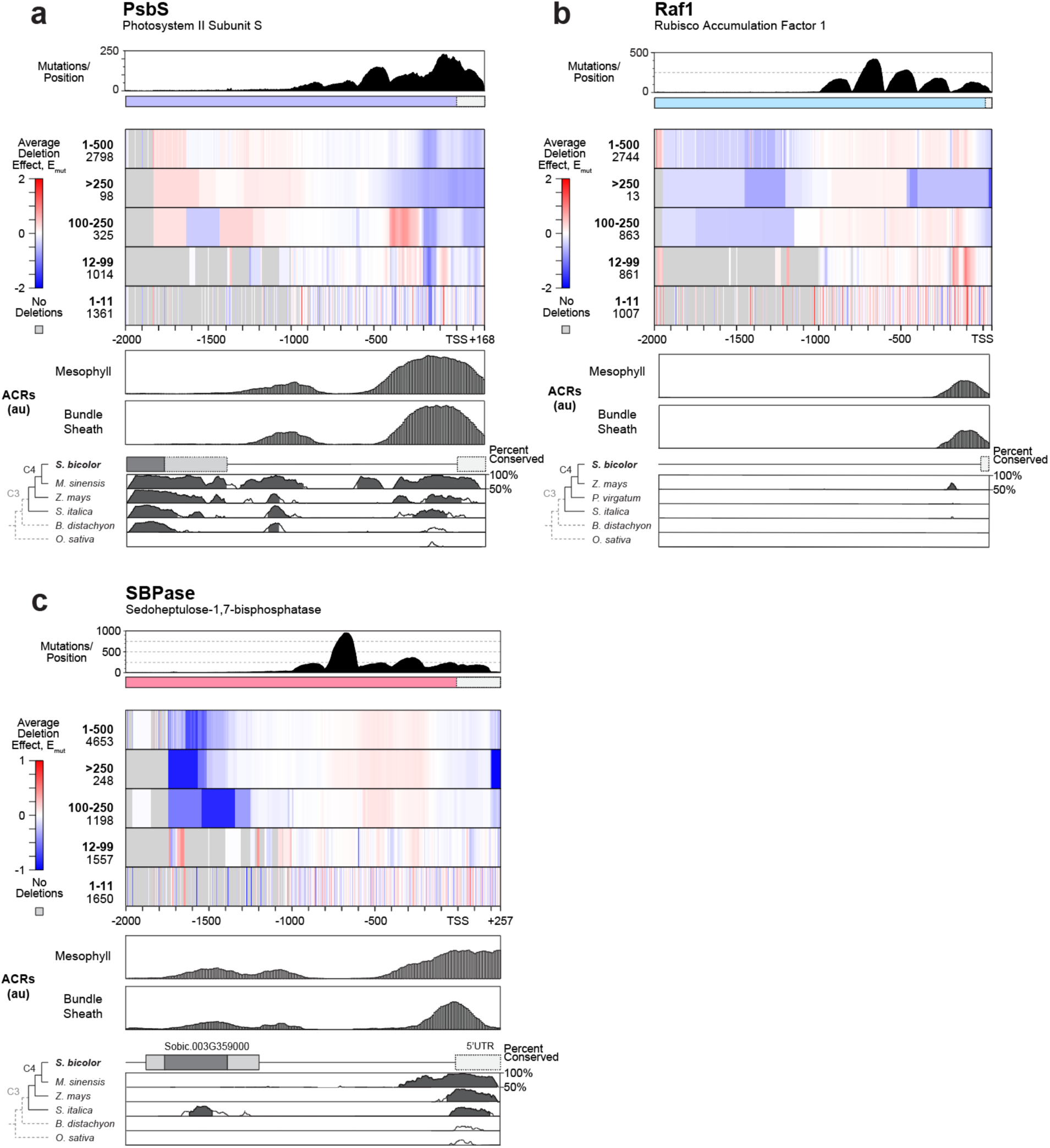
Deletion scanning across three photosynthetic genes. **a-c**, Histogram showing number of deletions and substitution per bp position, heatmap showing deletion outcomes in deletion size-denominated bins, accessible chromatin regions (ACRs) obtained from single-cell ATAC-seq (constant axes for all genes, arbitrary units)^36^, and local conservation to related C3 and C4 grasses for (**a**) *PsbS*, (**b**) *Raf1*, and (**c**) *SBPase*. For local conservation, sequence homology is plotted with 50% and 100% identity bounds, where shading indicates contiguous stretches >100 bp above 70% similarity.

**Supplementary Fig. 3.**
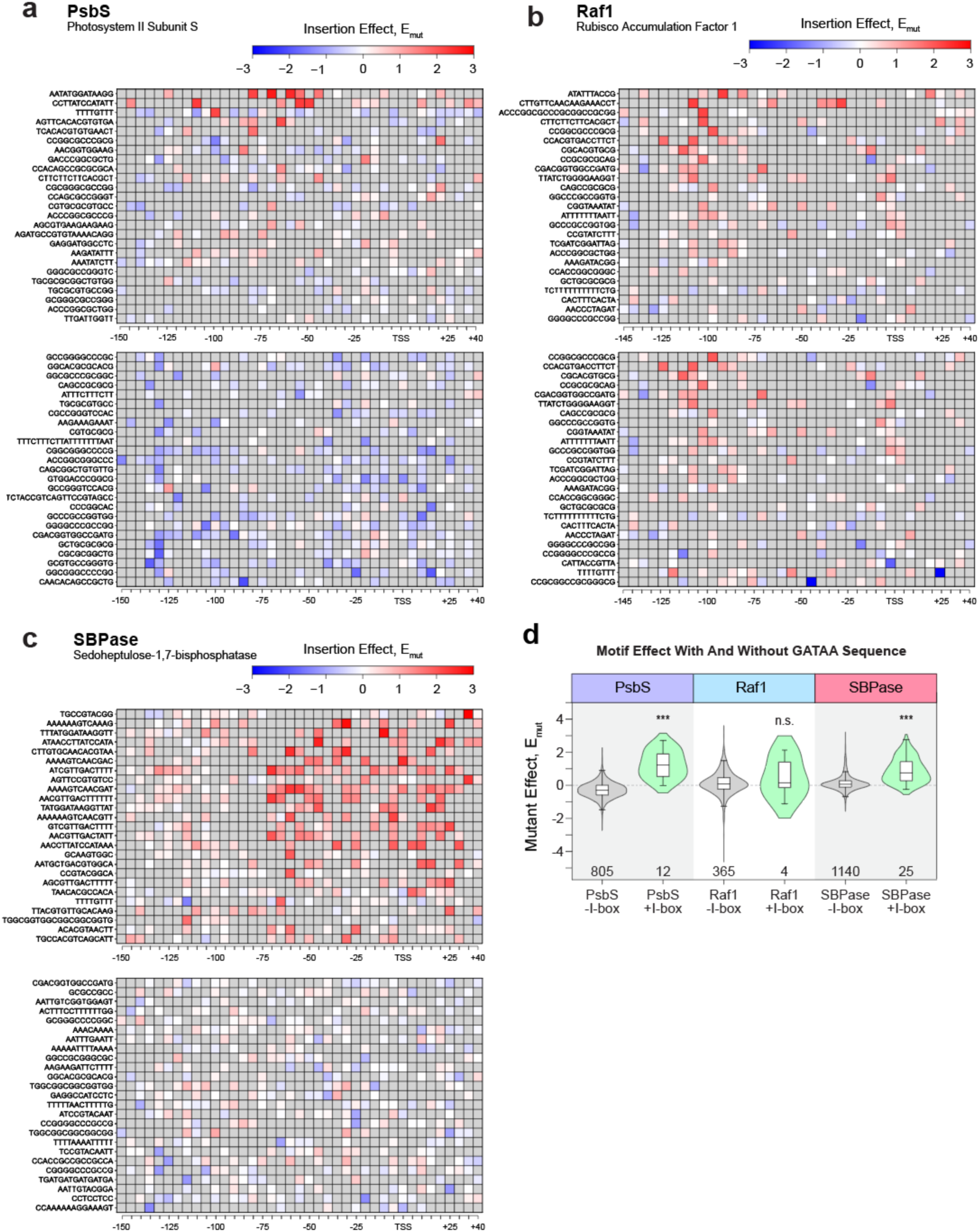
Activating and repressing motif insertions across genes. **a-c**, Insertion scanning across three photosynthetic genes reveals top 25 and bottom 25 enhancers ranked by maximum or minimum insertion effect for (**a**) PsbS, (**b**) Raf1, and (**c**) *SBPase*. **d**, Violin plot with overlaid box plots (center line, median; box limits, upper and lower quartiles; whiskers, 1.5× interquartile range) displaying the average effect of each motif insertion with (green) and without (grey) a GATAA/minimal I-box motif. P-values (asterisks) reflect Welch’s two-sided t-test in comparison to WT, with a permutation test to determine p-values (*** indicates p < 0.001).

